## Supporting information for "Exploring the repository of *de novo* designed bifunctional antimicrobial peptides through deep learning"

### Supplementary Figures

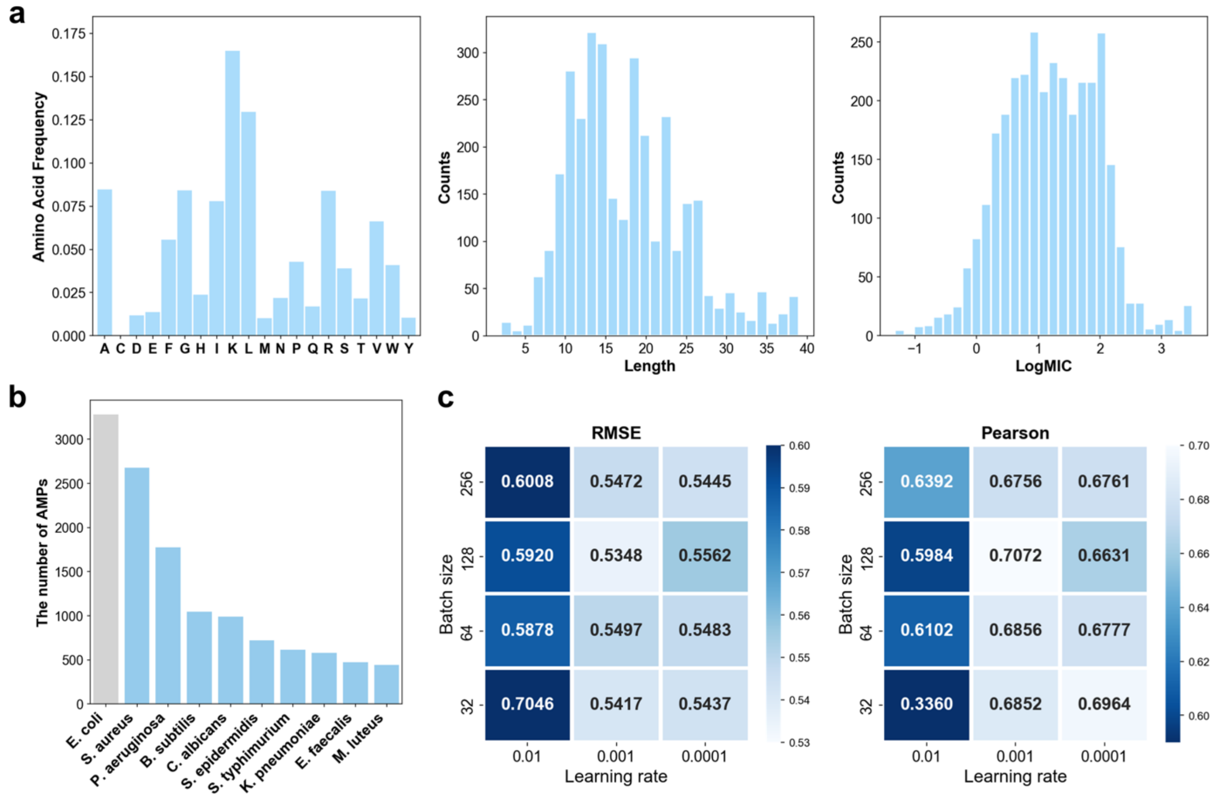


**Figure S1.** Dataset and hyperparameter settings of AMPredictor.

a. Distributions of amino acid compositions, sequence length, and antimicrobial activity numeric labels (logMIC).

b. Top ten kinds of microbes related to the AMPs in training set and the number of AMPs.

c. Grid search results of the batch size and learning rate, measured with RMSE and Pearson correlation coefficient.

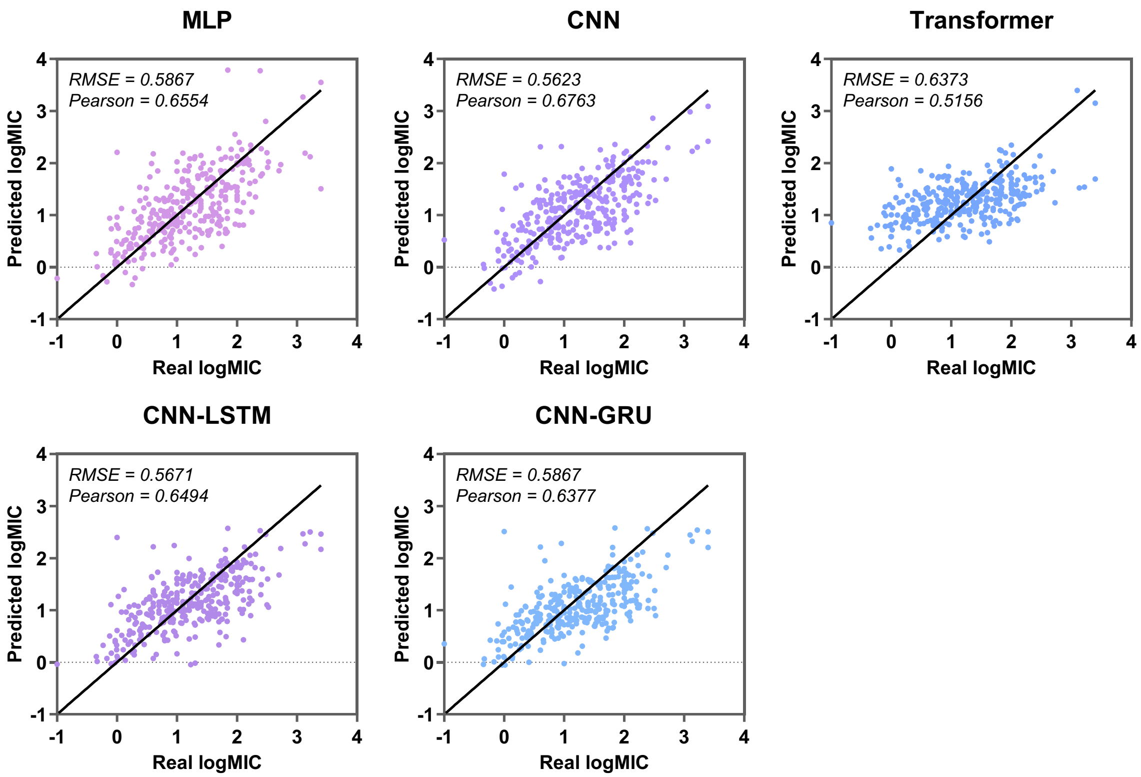


**Figure S2.** Antimicrobial activity regression results on test set of five baseline models.

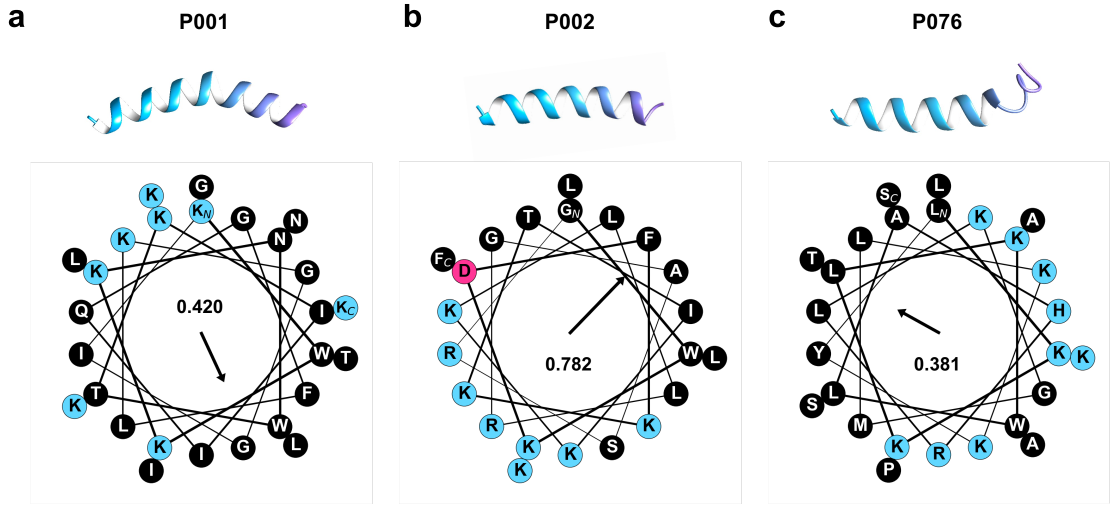


**Figure S3.** Predicted structures and the helical wheel projection of three selected peptides. Structures are predicted by AlphaFold2 (single sequence version). Helical wheels are plotted with modlAMP package and the Eisenberg hydrophobic moments are calculated and shown in the center.[1]

a. P001. b. P002. c. P076.

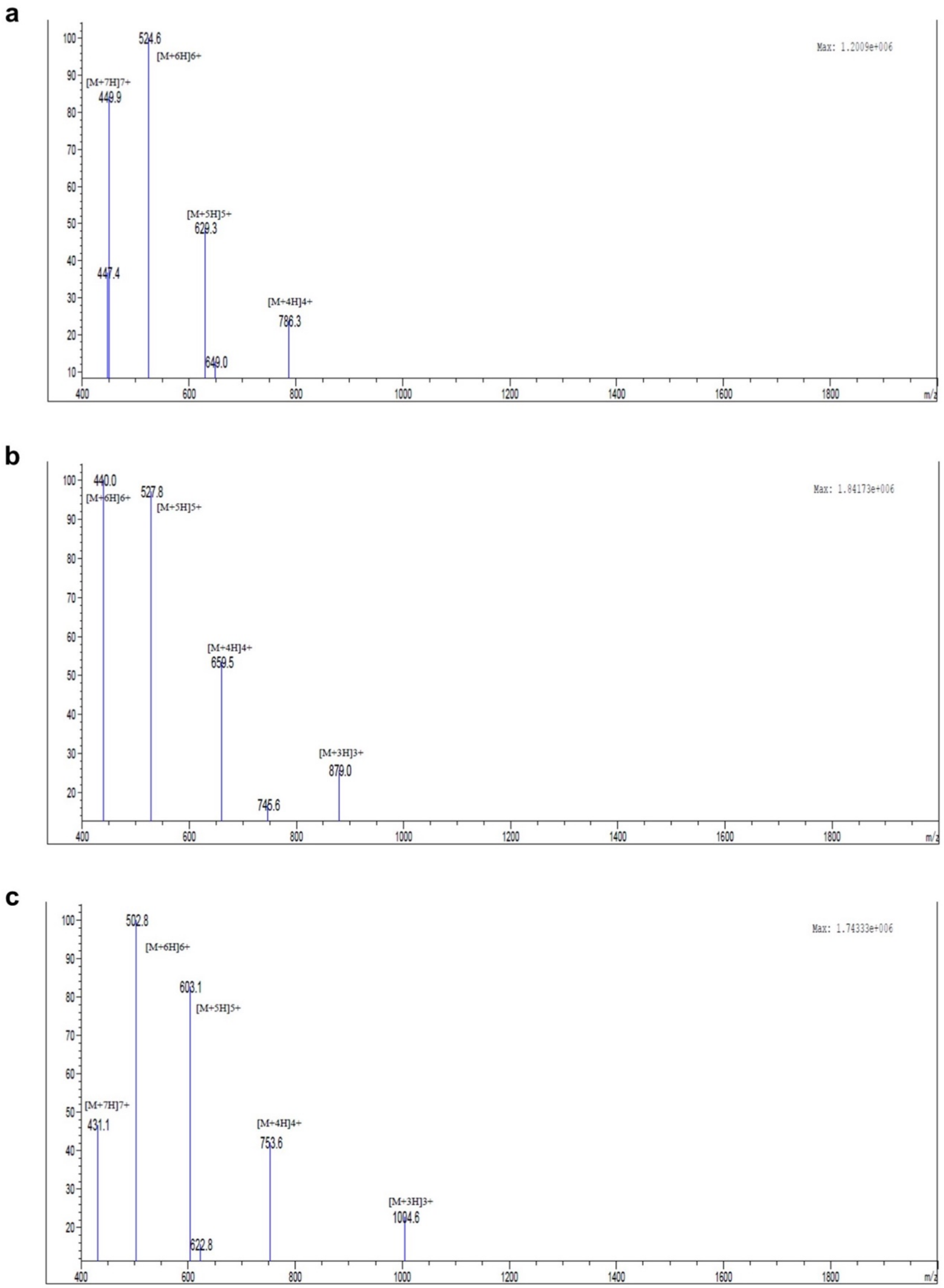


**Figure S4.** Mass spectrometry analysis. a. P001. b. P002. c. P076.

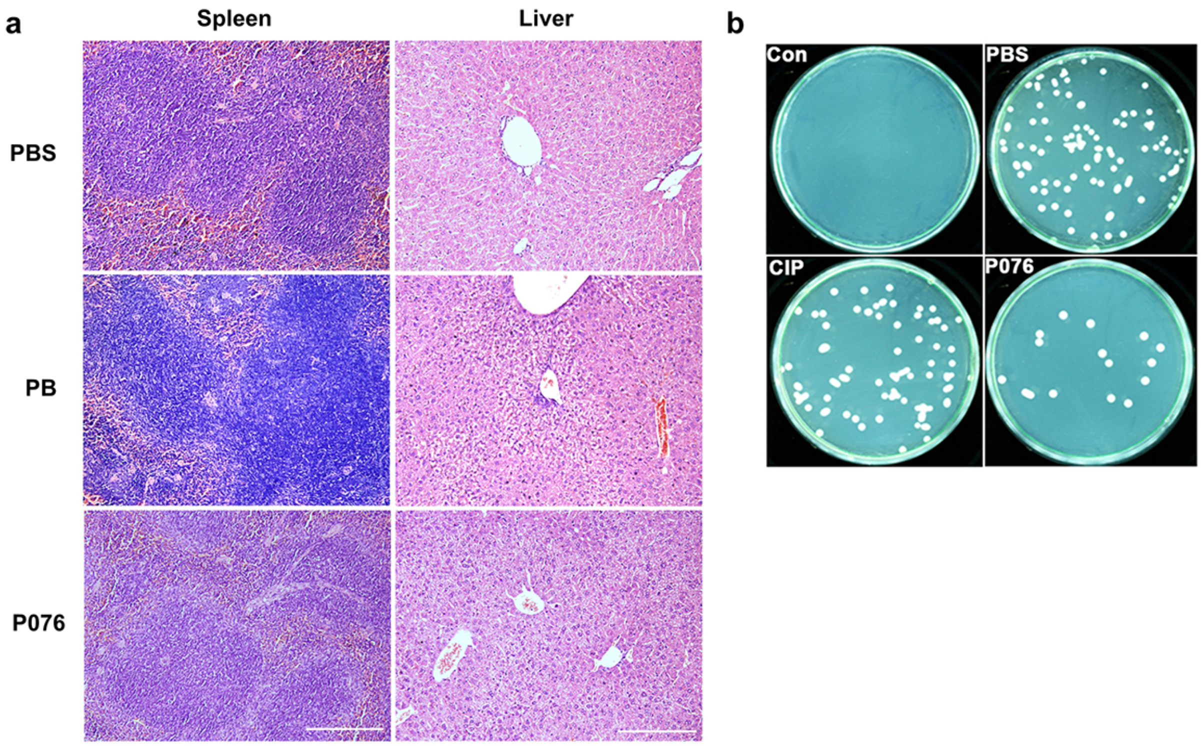


**Figure S5.** Additional results for in vivo antibacterial evaluations of P076.

a. H&E staining of mouse spleens and livers. The scale bar is 200 μm.

b. The bacterial colony on MHB plates with mouse spleen suspension.

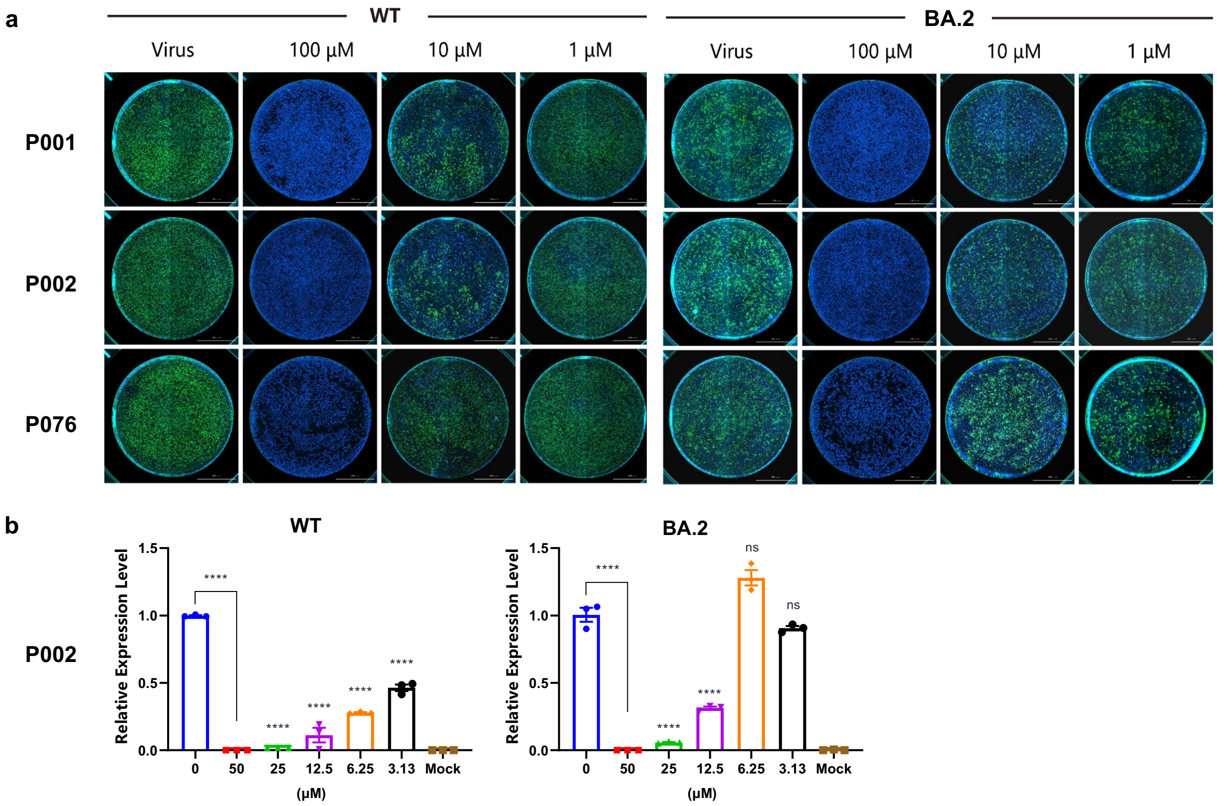


**Figure S6.** Antiviral assays of peptides against SARS-CoV-2 wild-type and BA.2 strains.

a. Immunofluorescence assays of P001, P002, and P076.

b. qRT-PCR of P002.

All experiments are performed in triplicates. Column bars are means ± SEMs. $^{****}$ P<0.0001. ns, no significance.

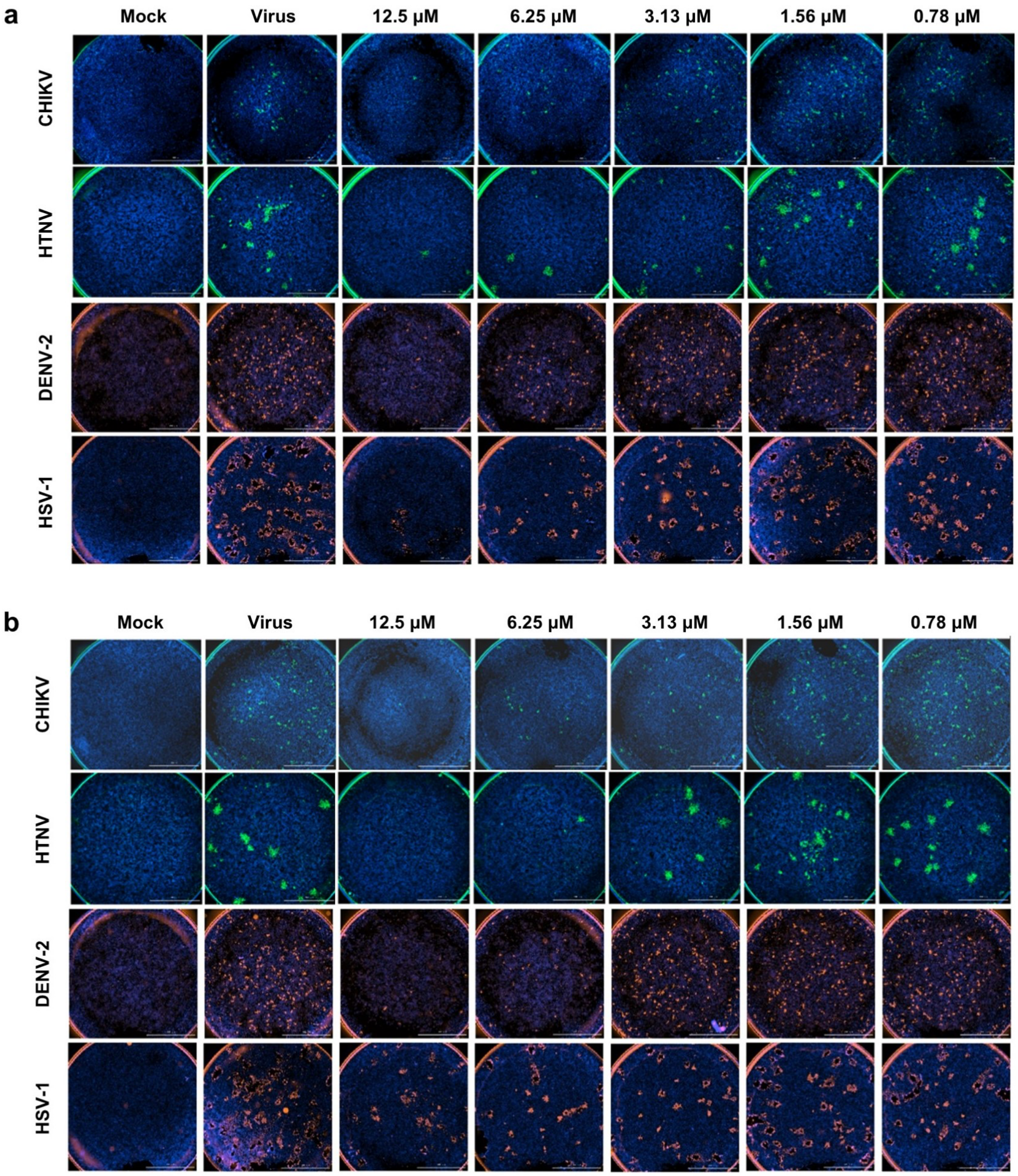


**Figure S7.** Immunofluorescence assays of P001 and P076 inhibiting CHIKV, HTNV, DENV-2, and HSV-1. All experiments are performed in triplicates.

a. P001. b. P076.

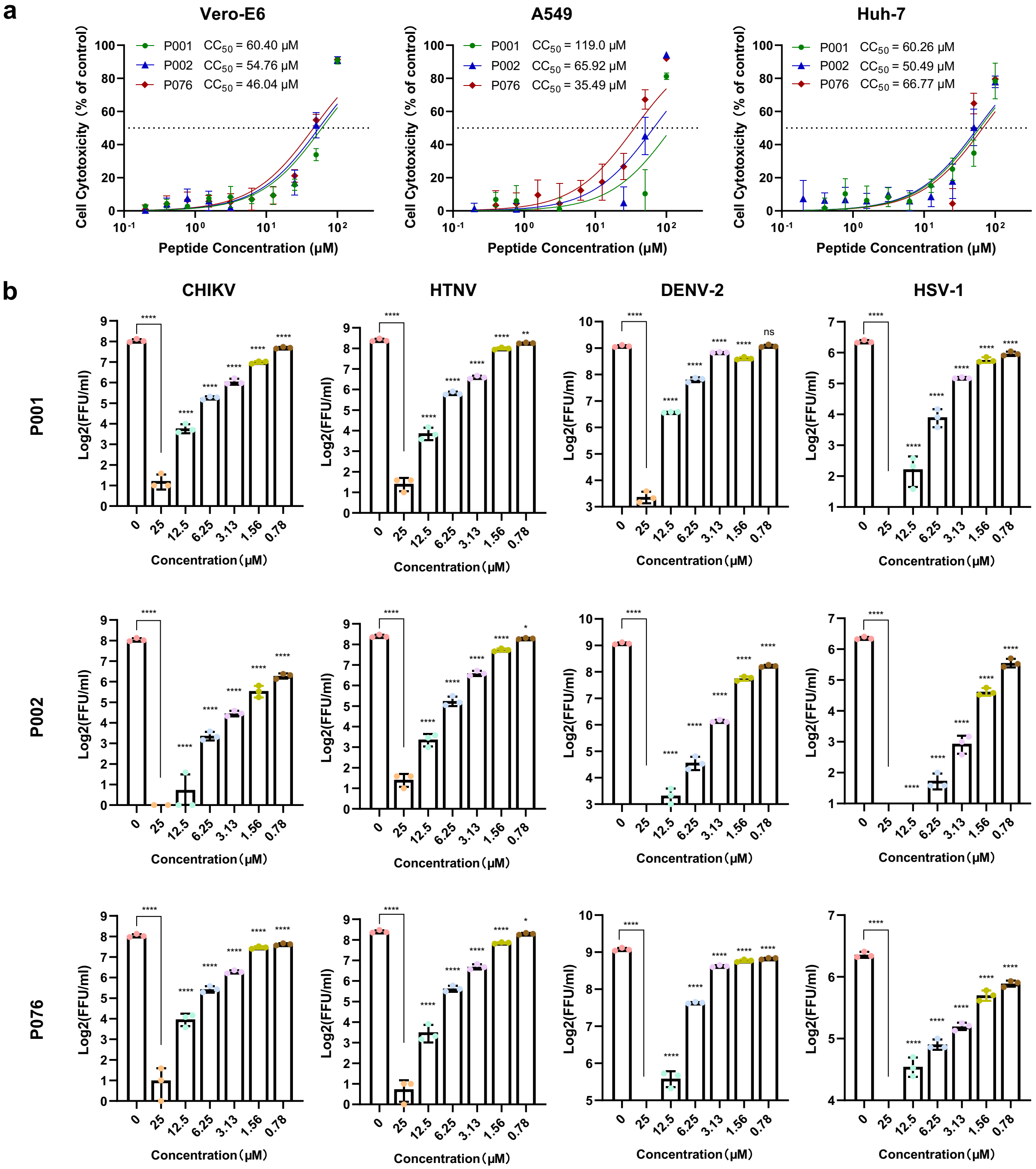


**Figure S8.** Cytotoxicity and quantitative immunofluorescence results.

a. Cytotoxicity curve of P001, P002, and P076 against Vero-E6, A549, and Huh-7 cells.

b. Quantification of immunofluorescence assay for three AMPs against four viruses.

All experiments are performed in triplicates. Column bars are means ± SEMs. * P<0.05, ** P<0.01, *** P<0.001, **** P<0.0001. ns, no significance.

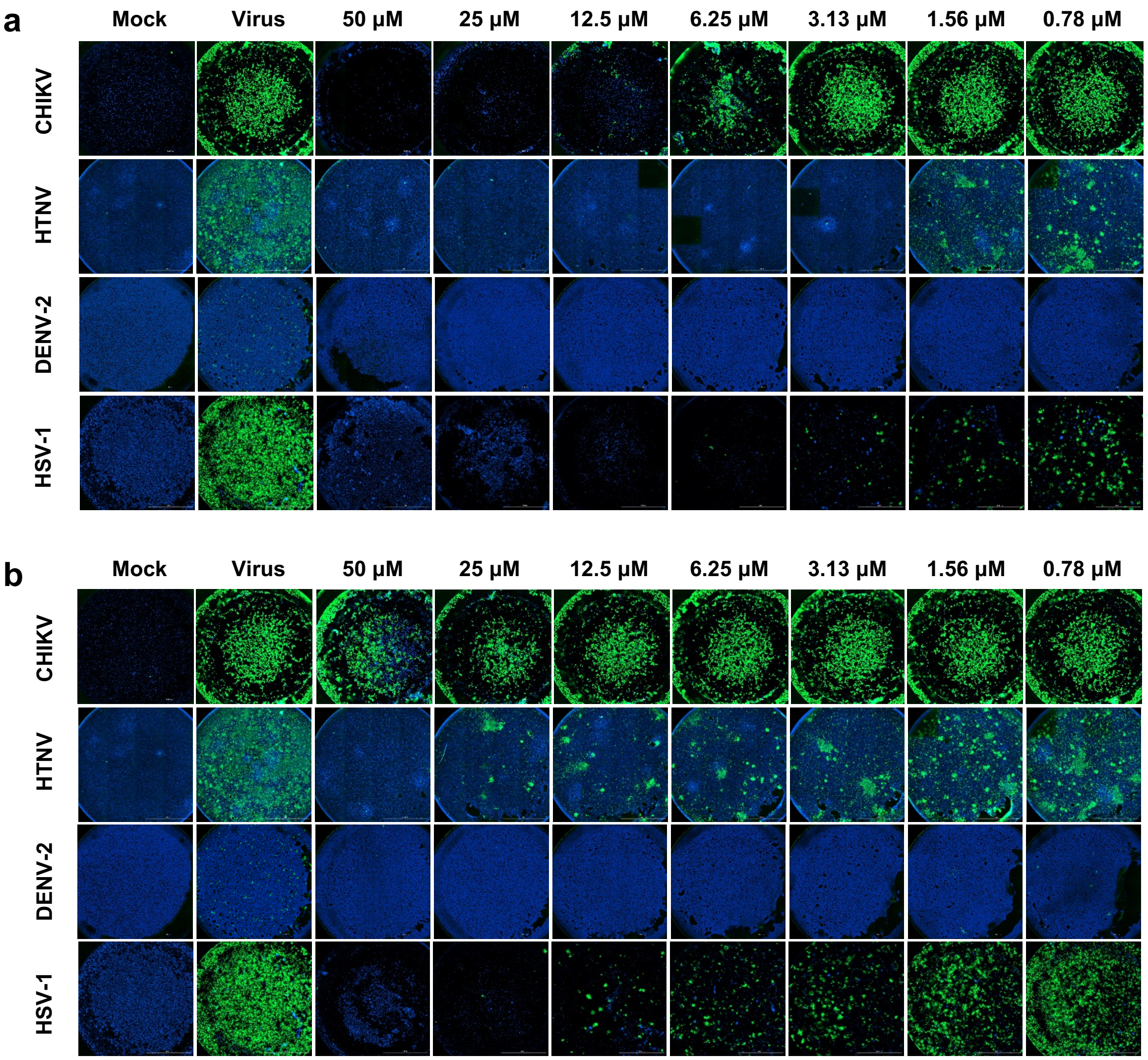


**Figure S9.** Immunofluorescence assays of P135 and P244 inhibiting CHIKV, HTNV, DENV-2, and HSV-1. All experiments are performed in triplicates.

a. P135. b. P244.

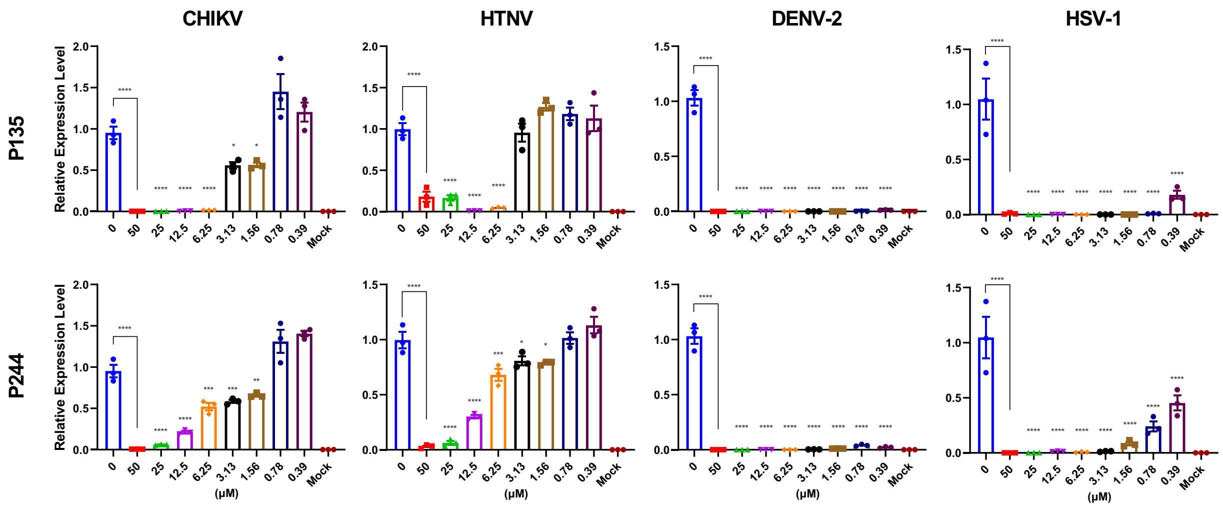


**Figure S10.** Quantitative real-time PCR of virus RNA with gradient concentrations of P135 and P244 inhibiting CHIKV, HTNV, DENV-2, and HSV-1. All experiments are performed in triplicates. * P<0.05, ** P<0.01, *** P<0.001, **** P<0.0001. ns, no significance.

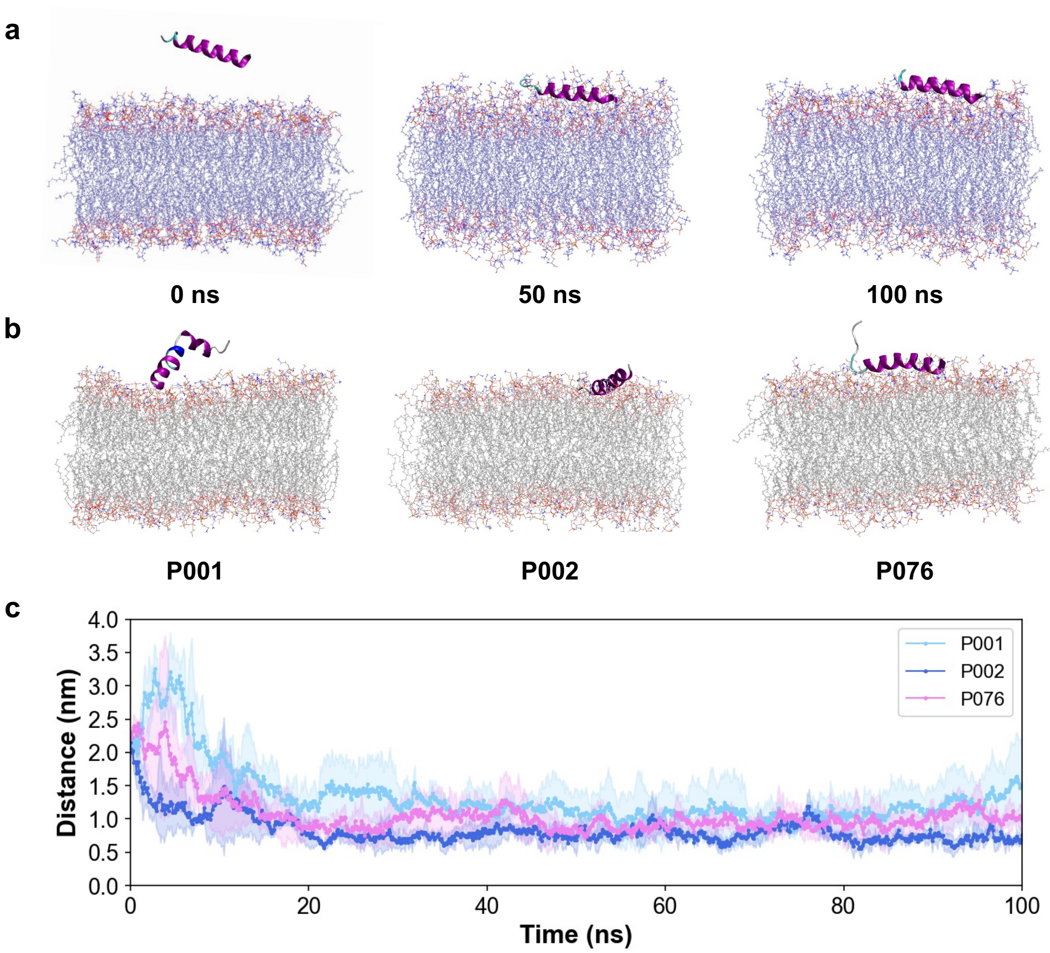


**Figure S11.** Molecular dynamics simulations for peptides and different lipid bilayers.

a. Snapshots of P002 with viral envelopes at 0 ns, 50 ns, and 100 ns.

b. Snapshots of P001, P002 and P076 with G- inner membranes at 100 ns.

c. Distance between the peptides and the viral membrane surface in the trajectories.

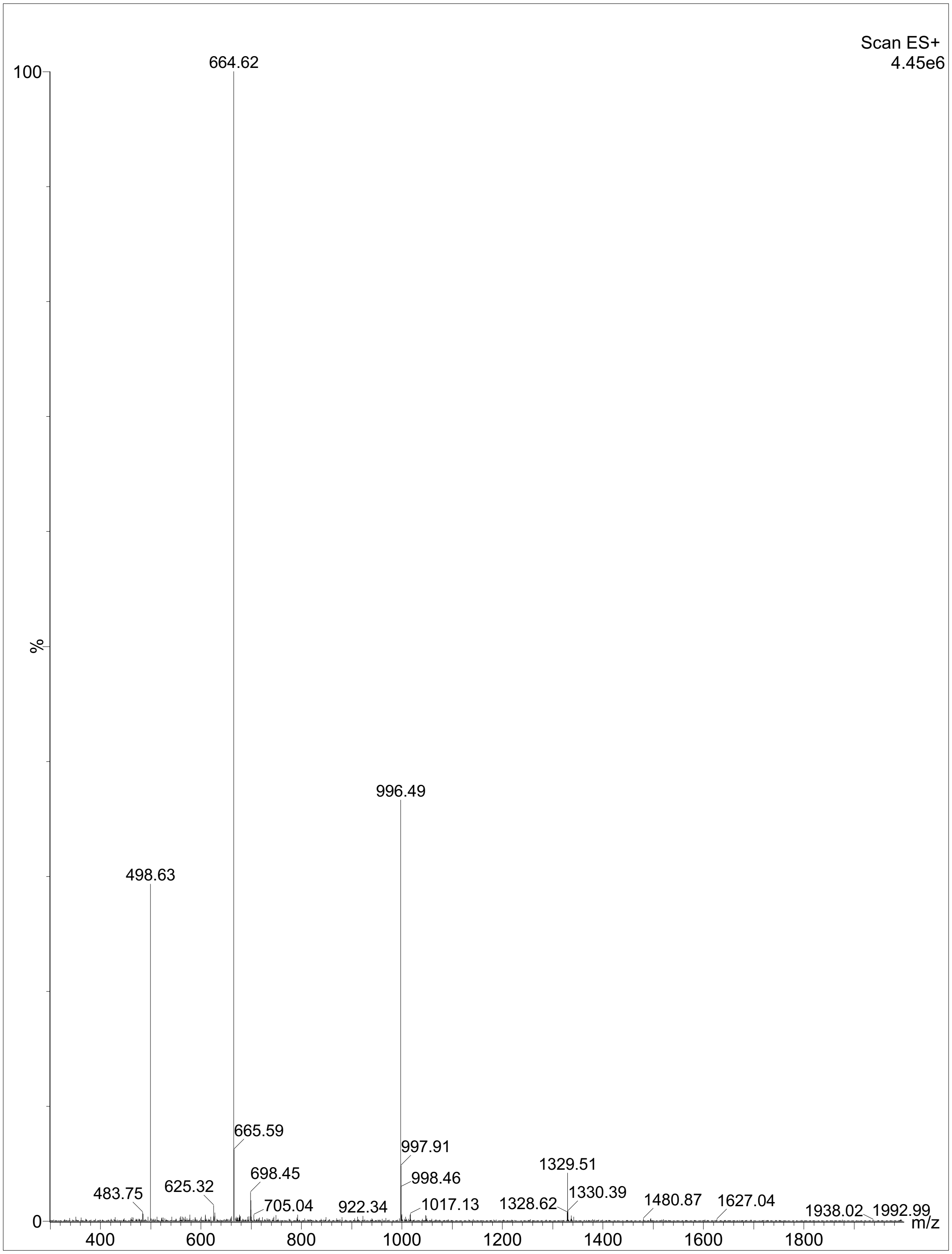


**Figure S12.** Mass spectrometry of P089.

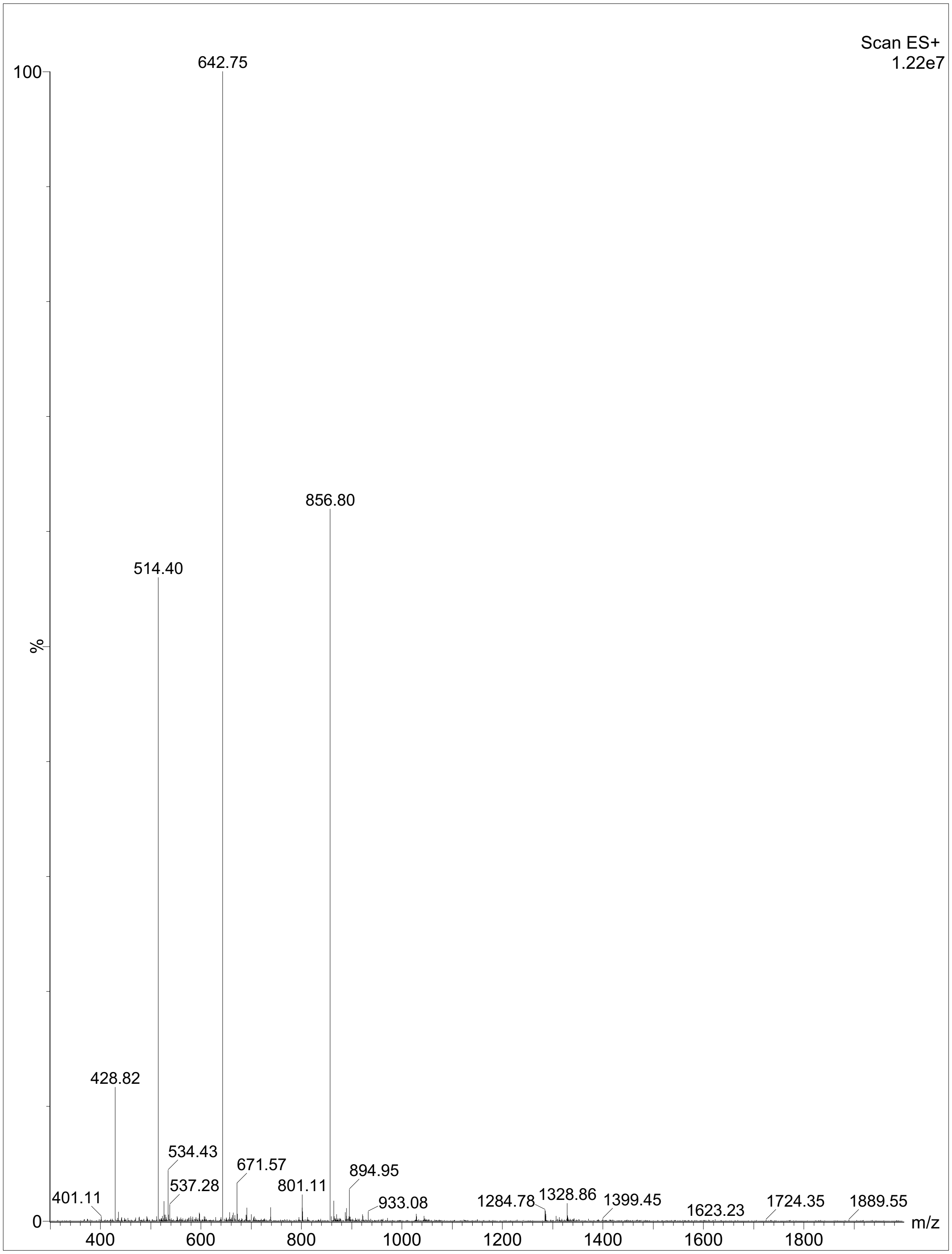


**Figure S13.** Mass spectrometry of P386.

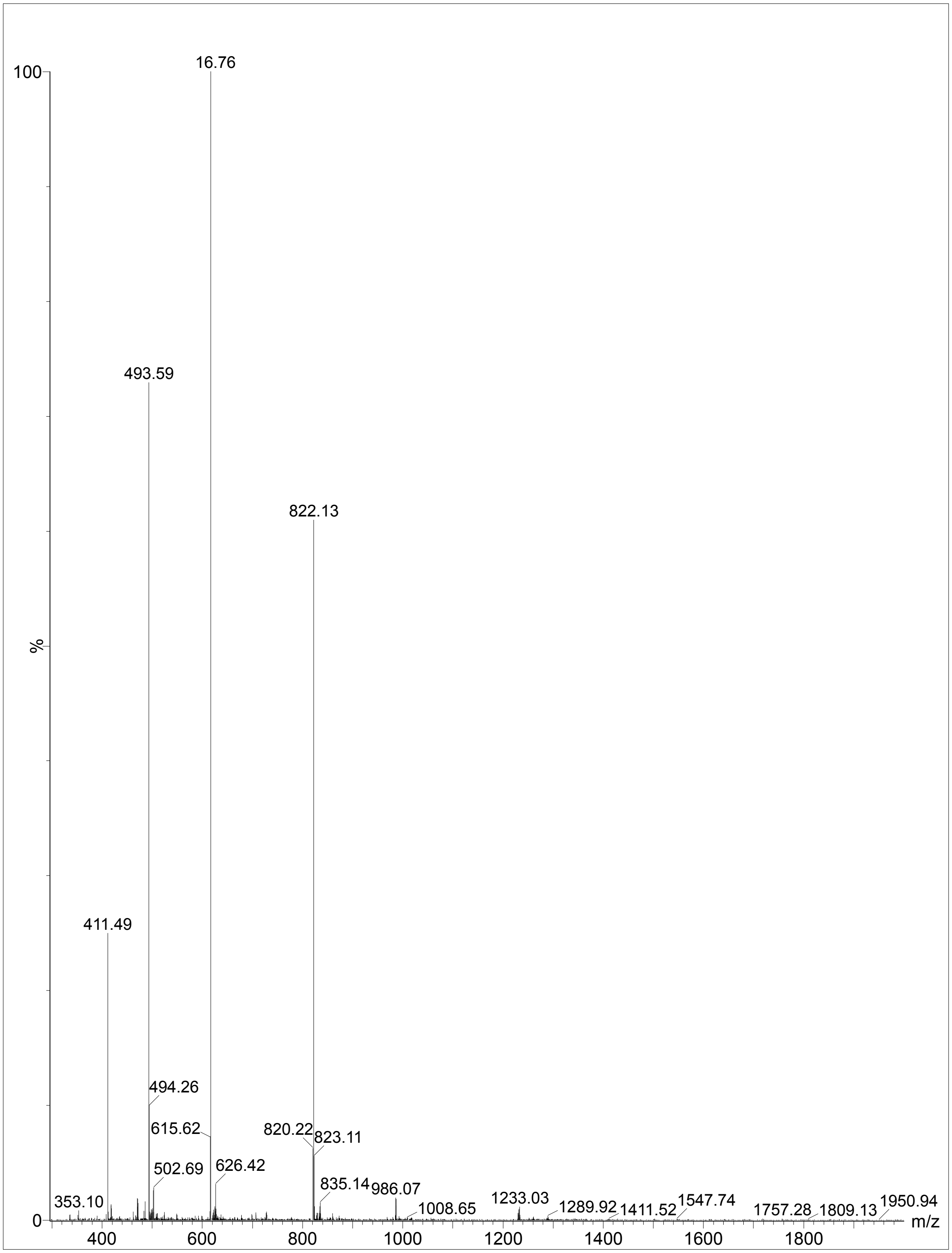


**Figure S14.** Mass spectrometry of P120.

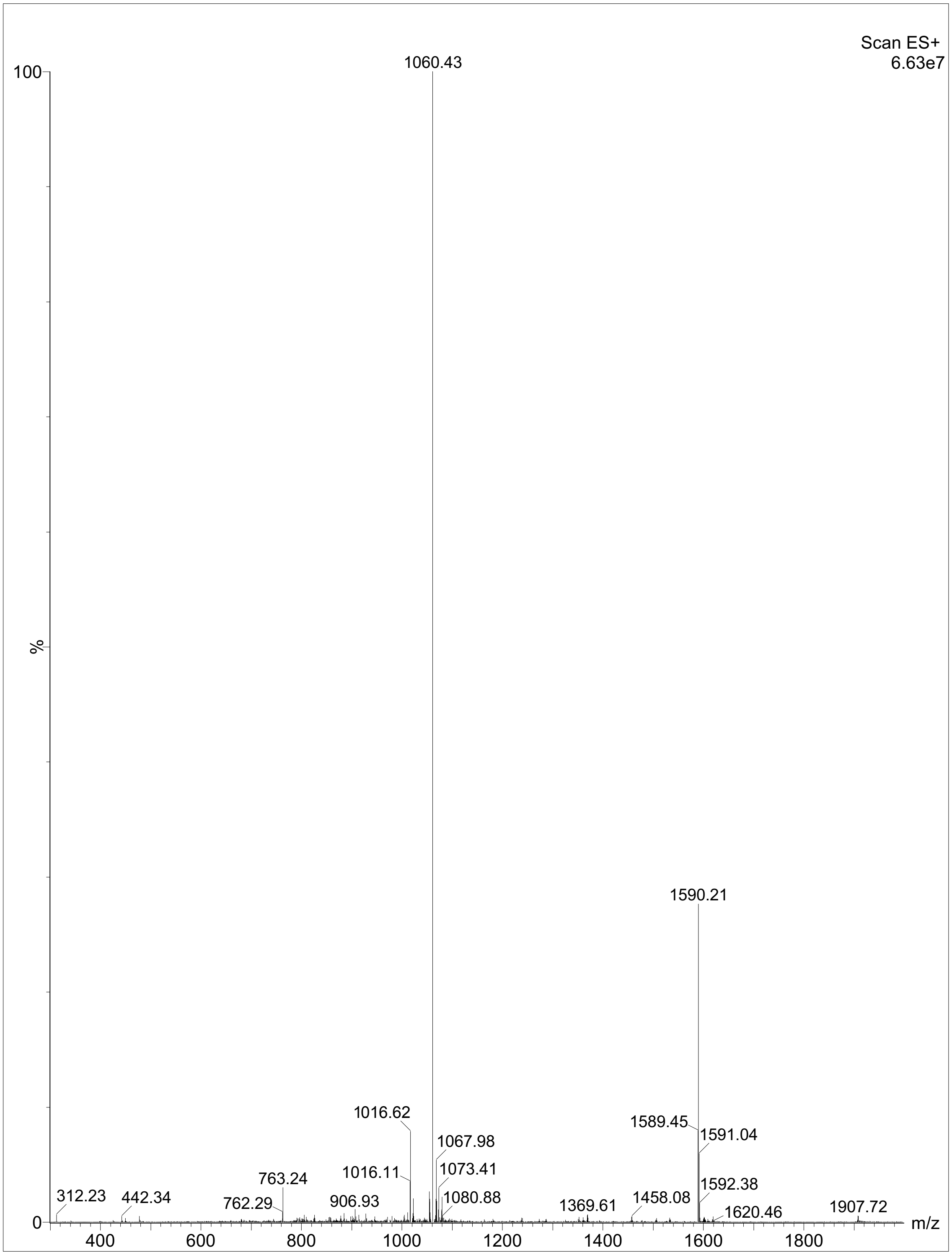


**Figure S15.** Mass spectrometry of P135.

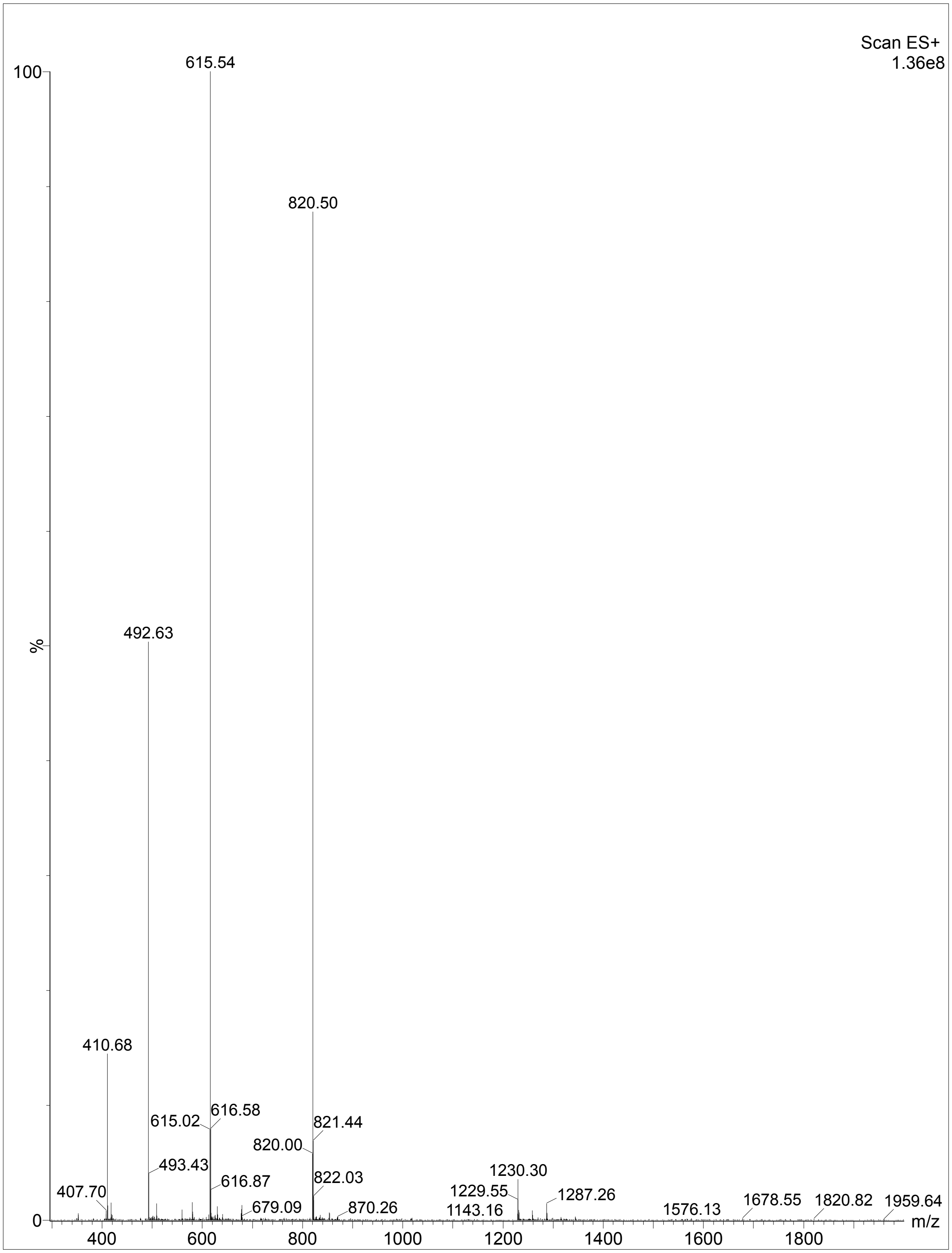


**Figure S16.** Mass spectrometry of P247.

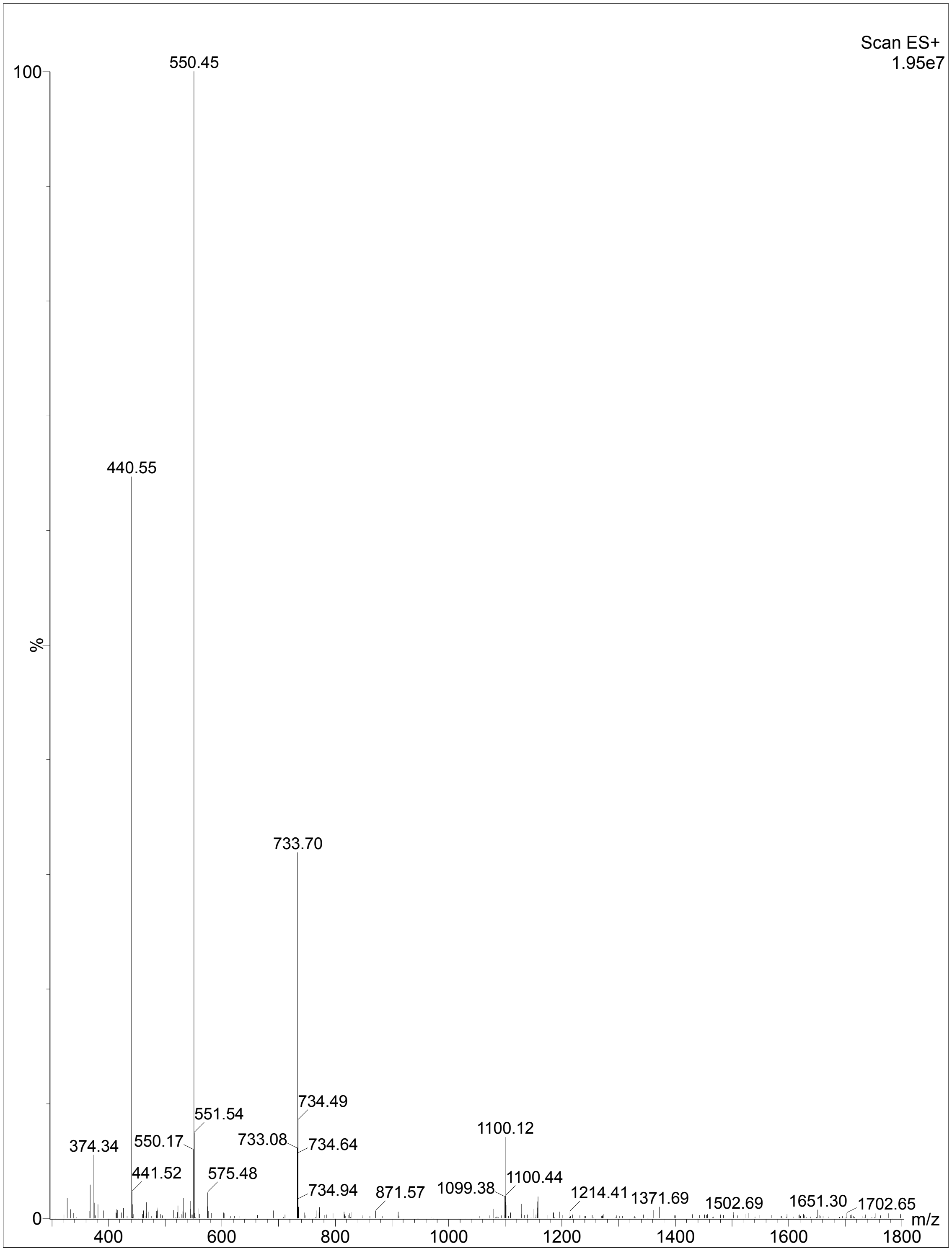


**Figure S17.** Mass spectrometry of P026.

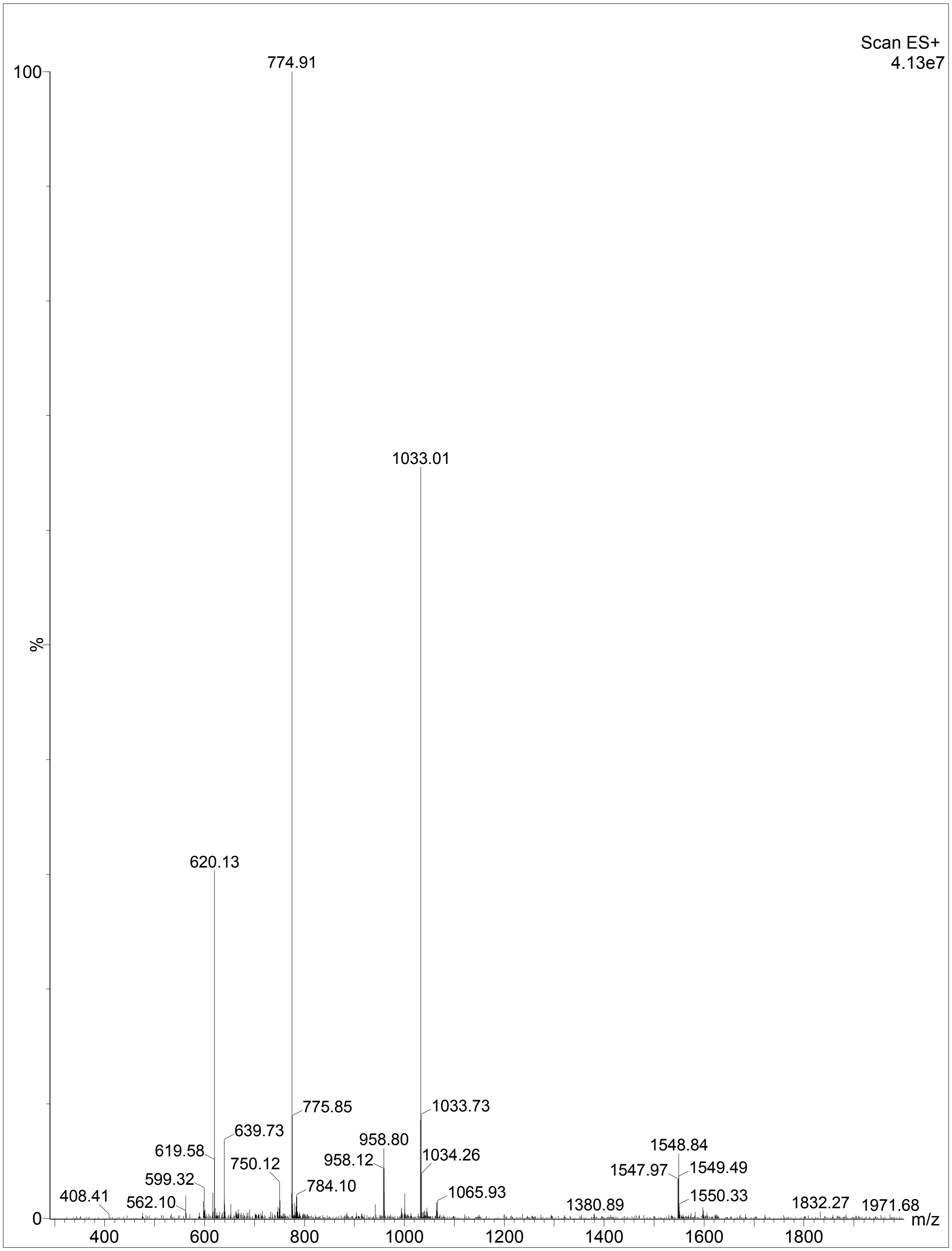


**Figure S18.** Mass spectrometry of P039.

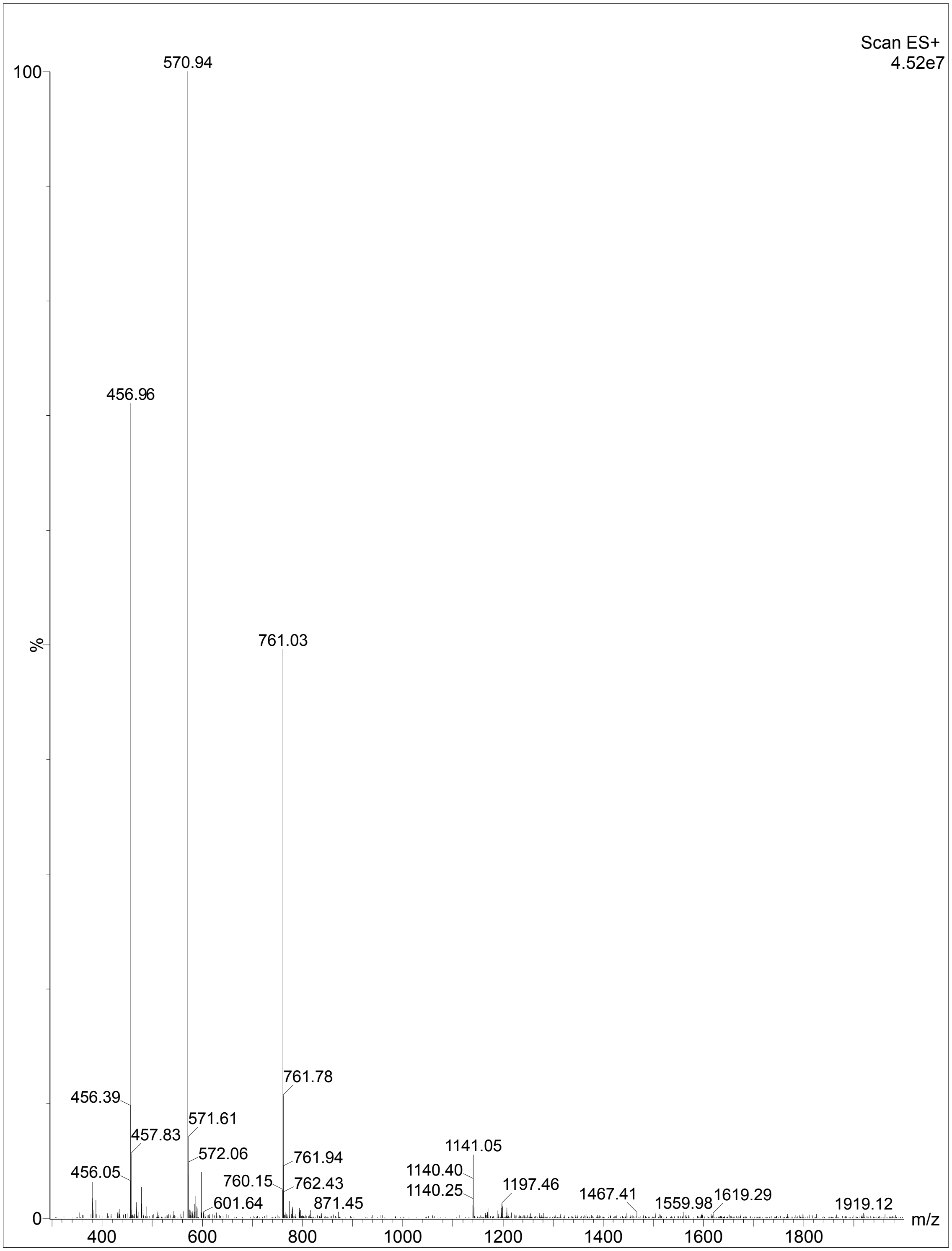


**Figure S19.** Mass spectrometry of P070.

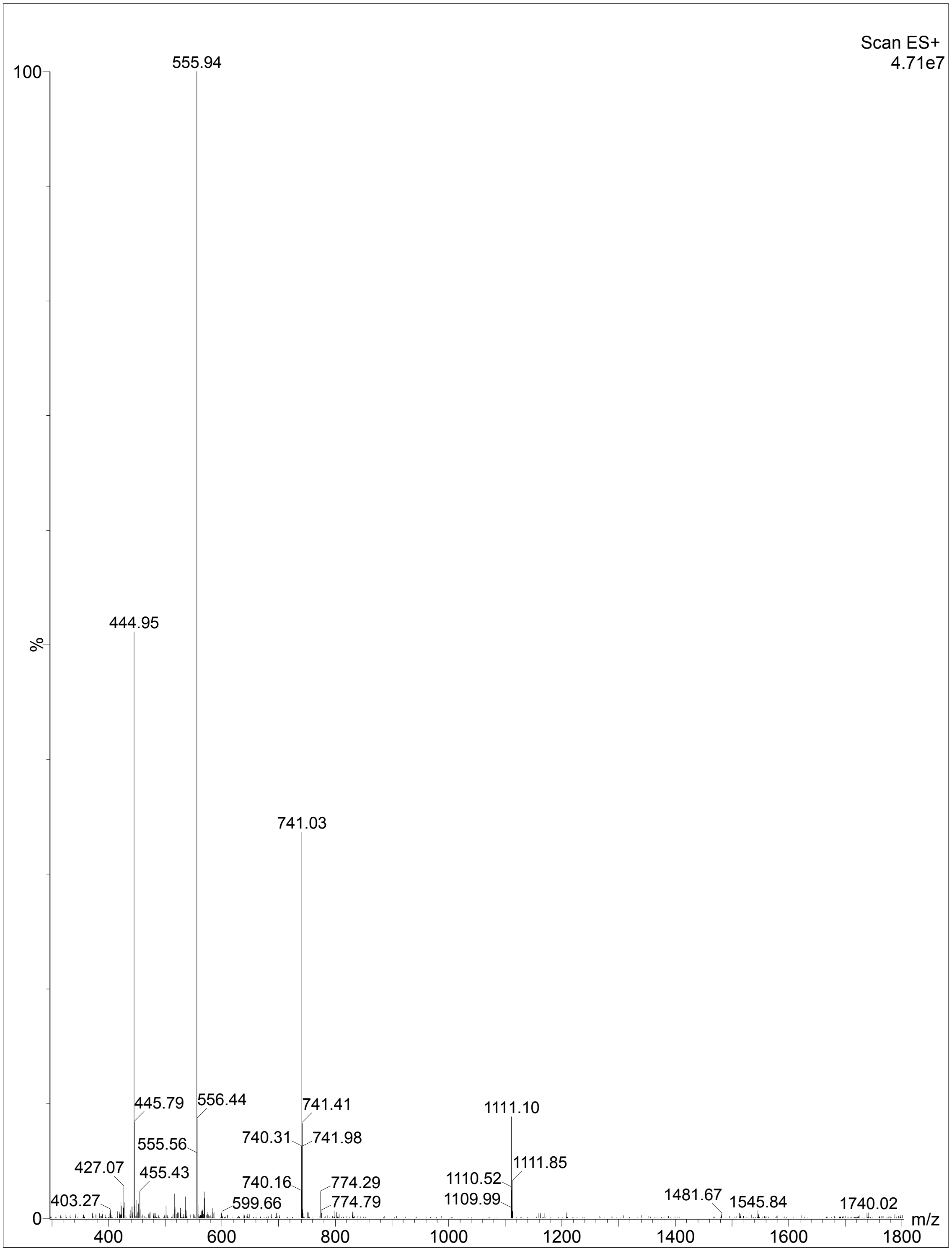


**Figure S20.** Mass spectrometry of P091.

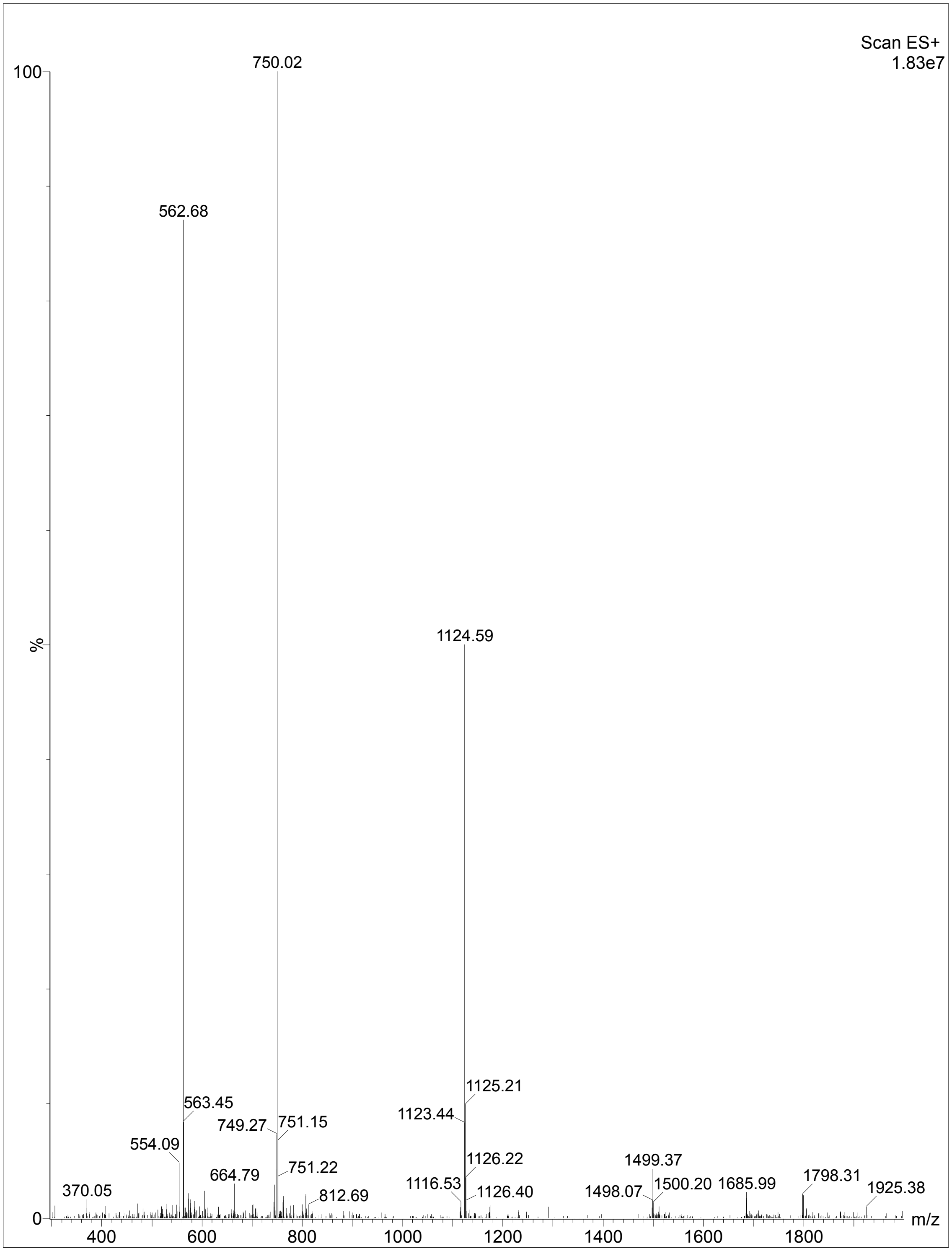


**Figure S21.** Mass spectrometry of P127.

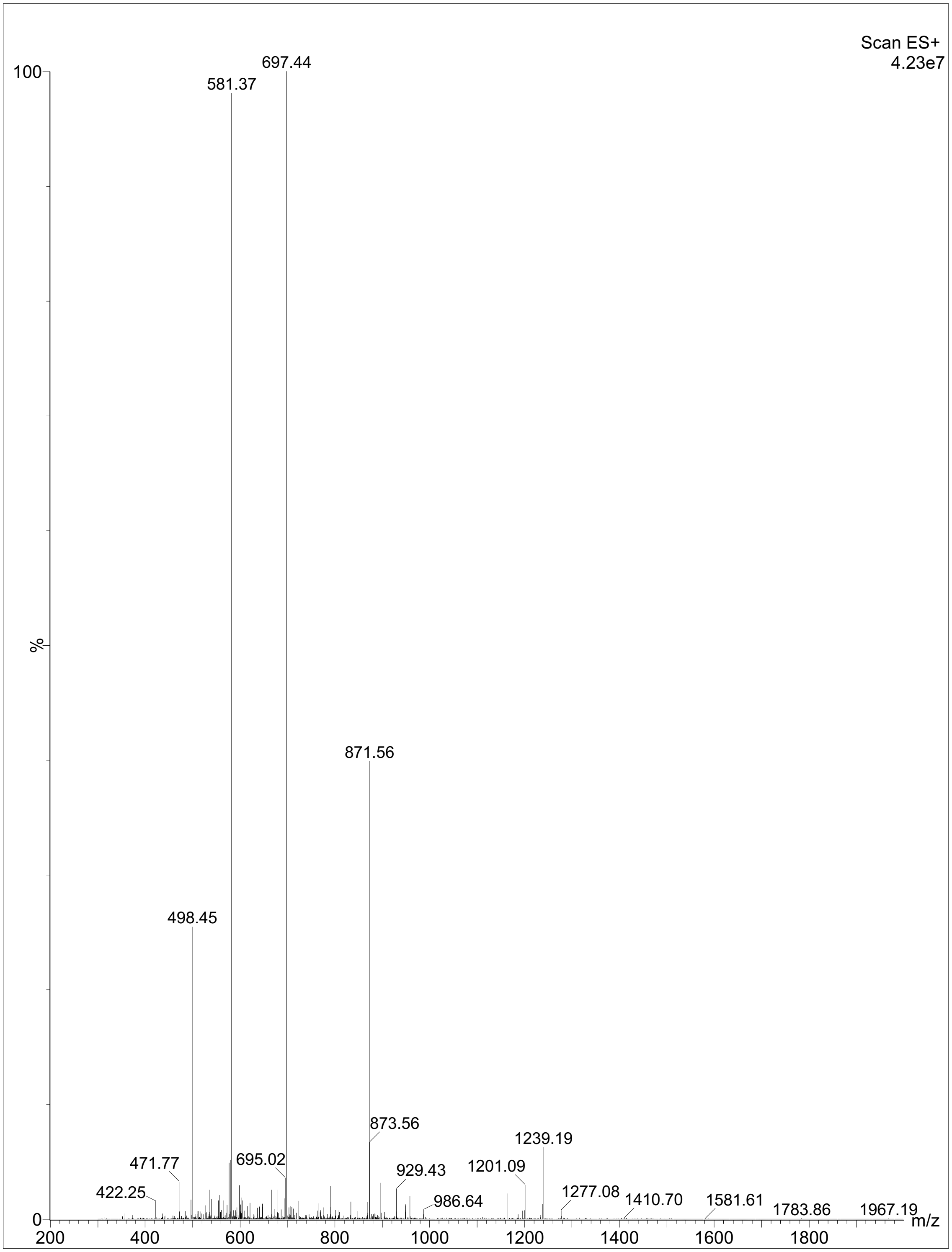


**Figure S22.** Mass spectrometry of P122.

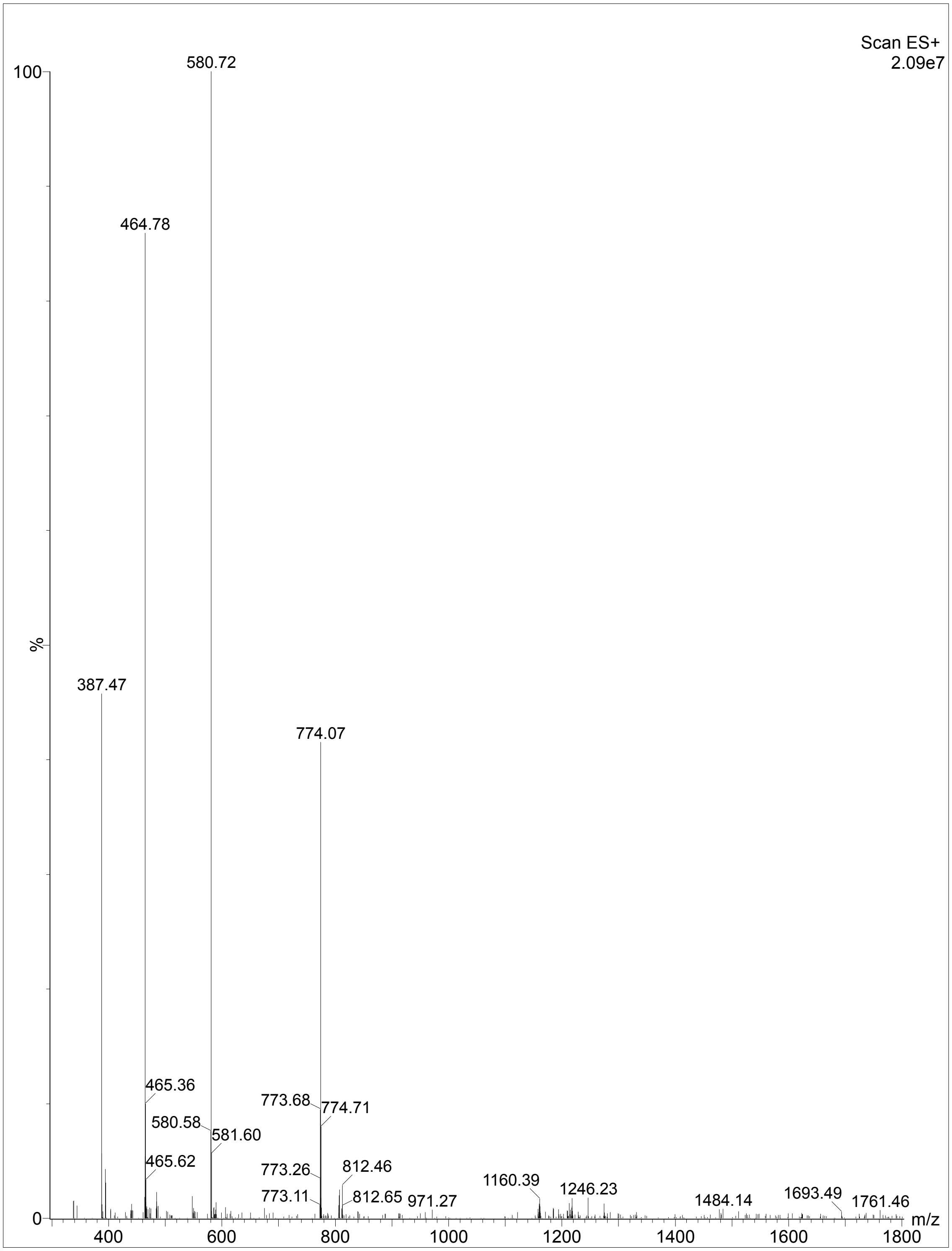


**Figure S23.** Mass spectrometry of P352.

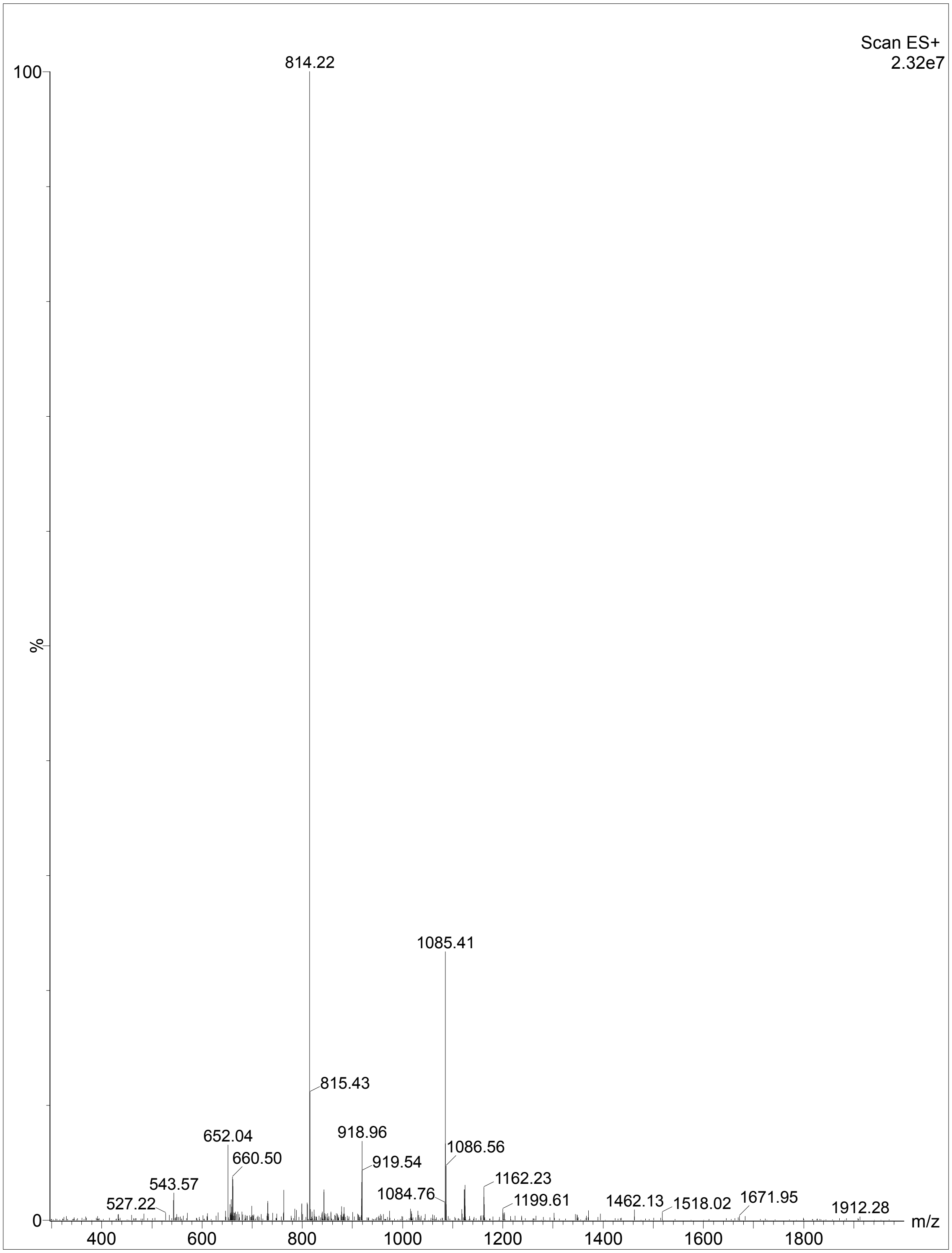


**Figure S24.** Mass spectrometry of P105.

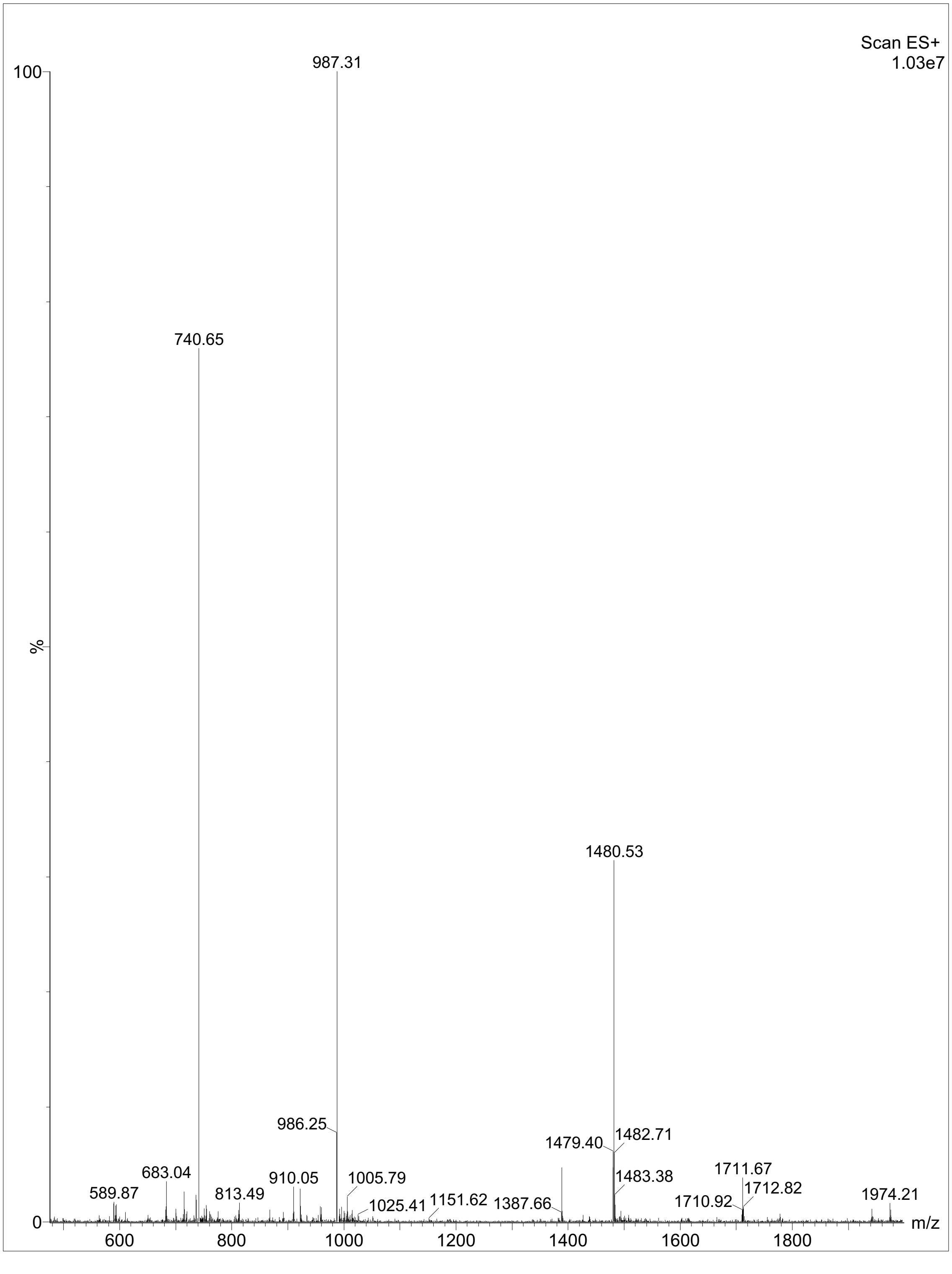


**Figure S25.** Mass spectrometry of P316.

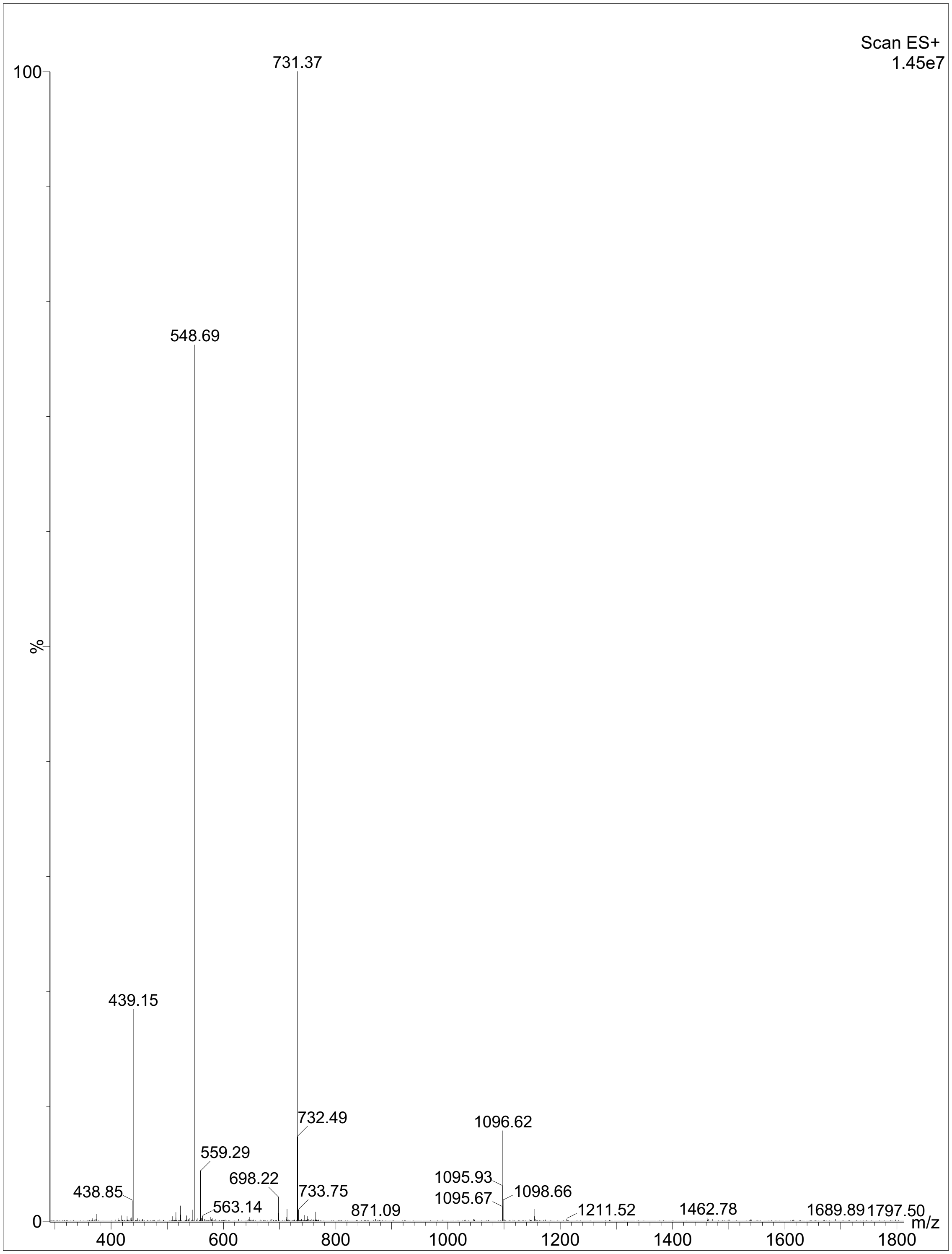


**Figure S26.** Mass spectrometry of P244.

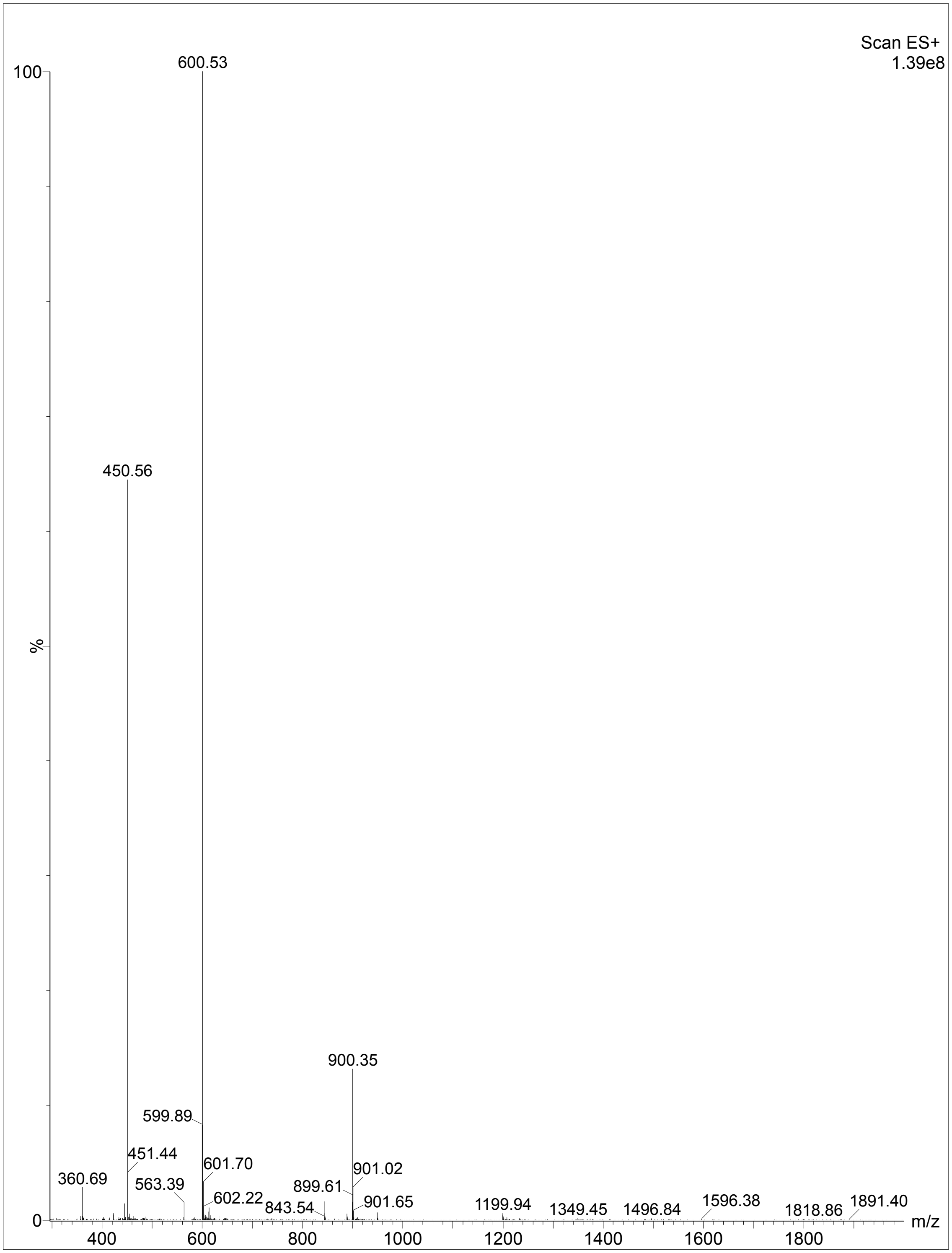


**Figure S27.** Mass spectrometry of P302.

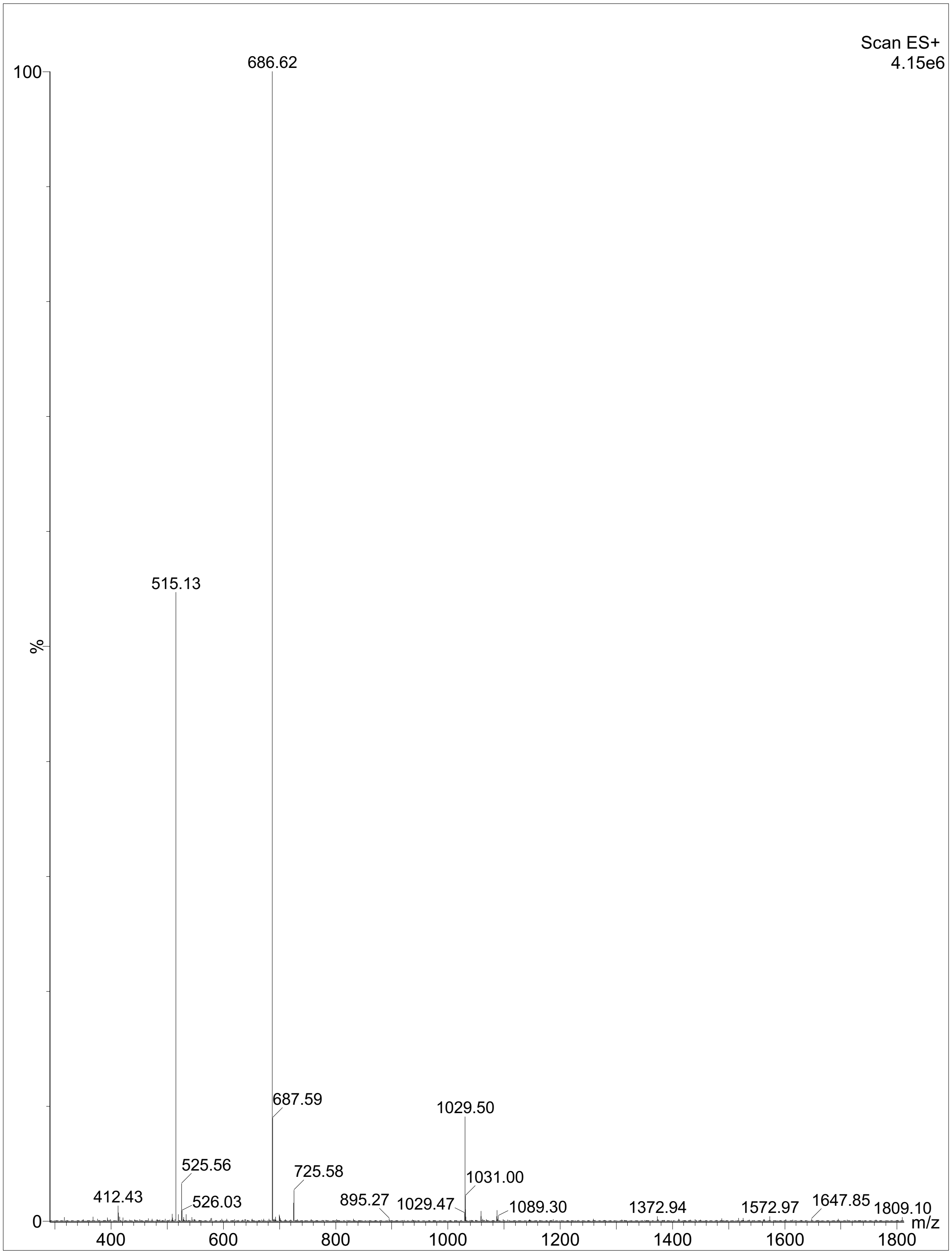


**Figure S28.** Mass spectrometry of P185.

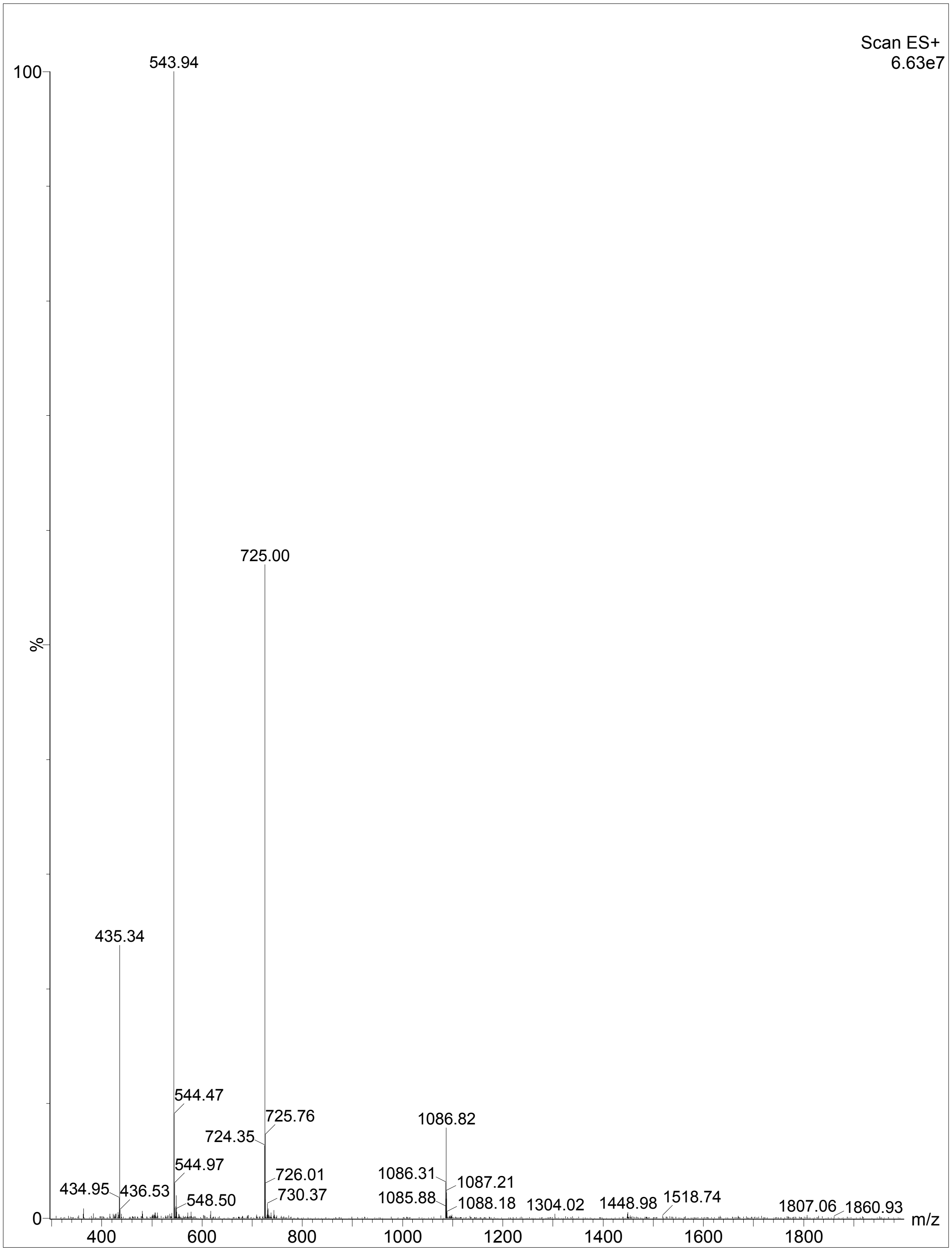


**Figure S29.** Mass spectrometry of P019.

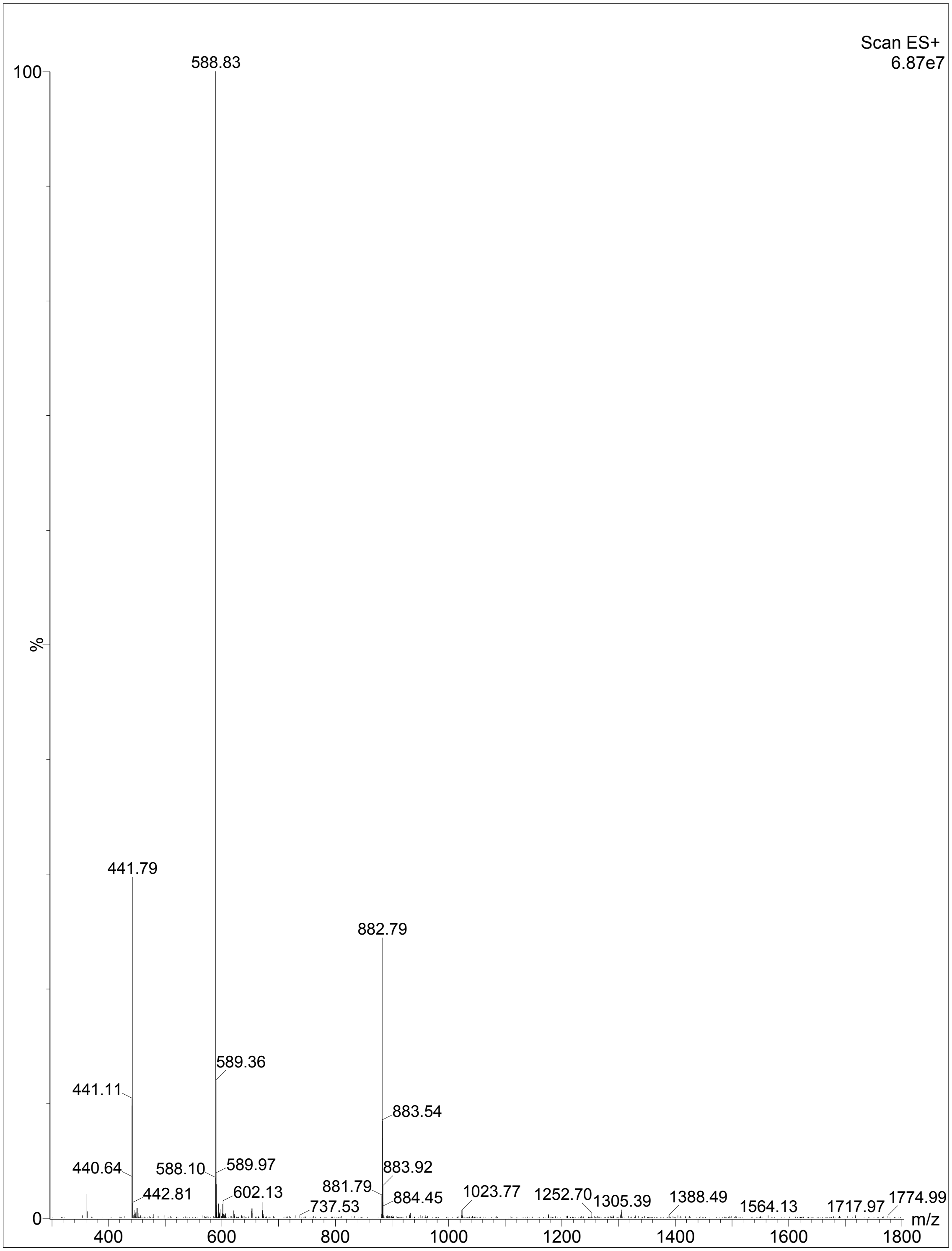


**Figure S30.** Mass spectrometry of P252.

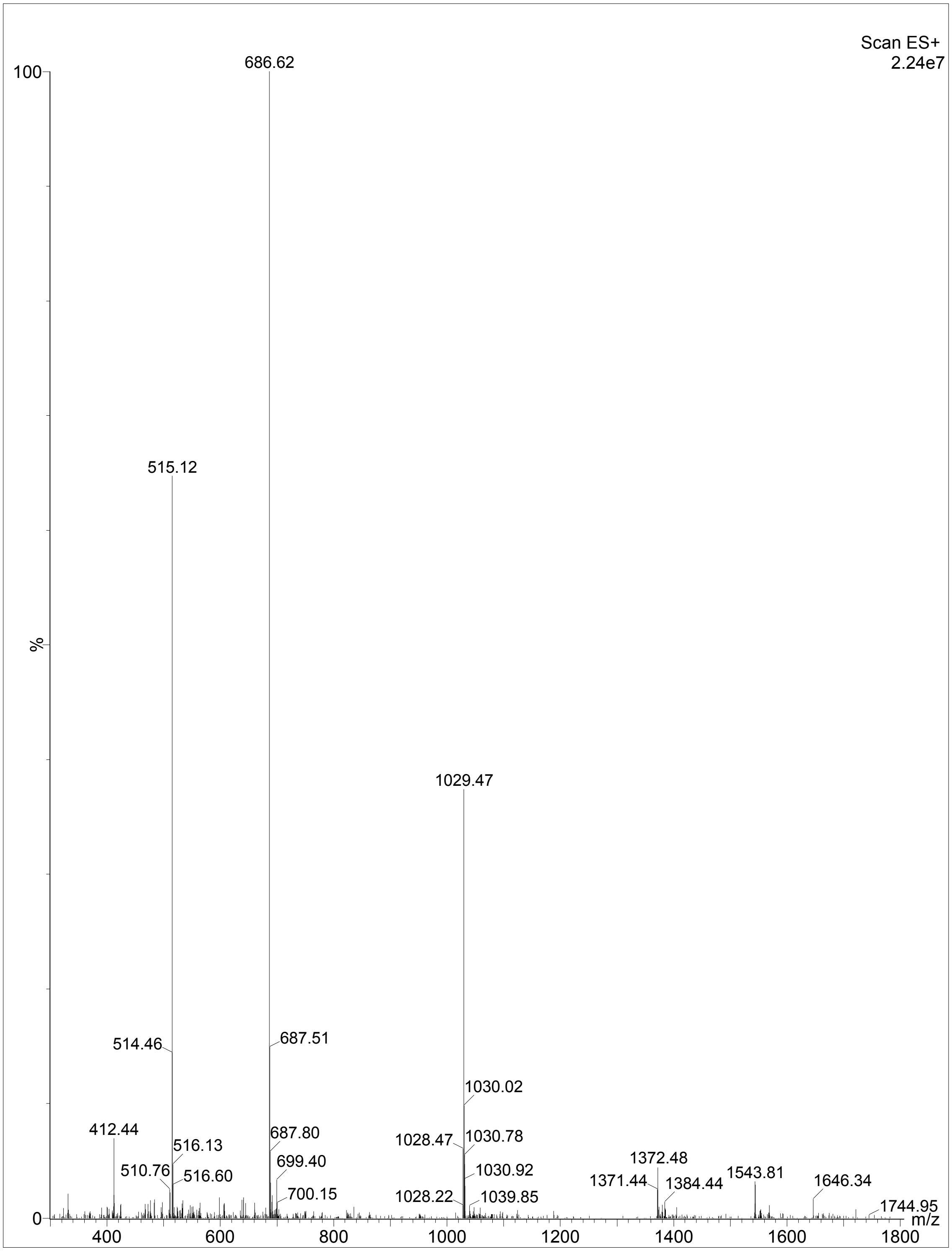


**Figure S31.** Mass spectrometry of P213.

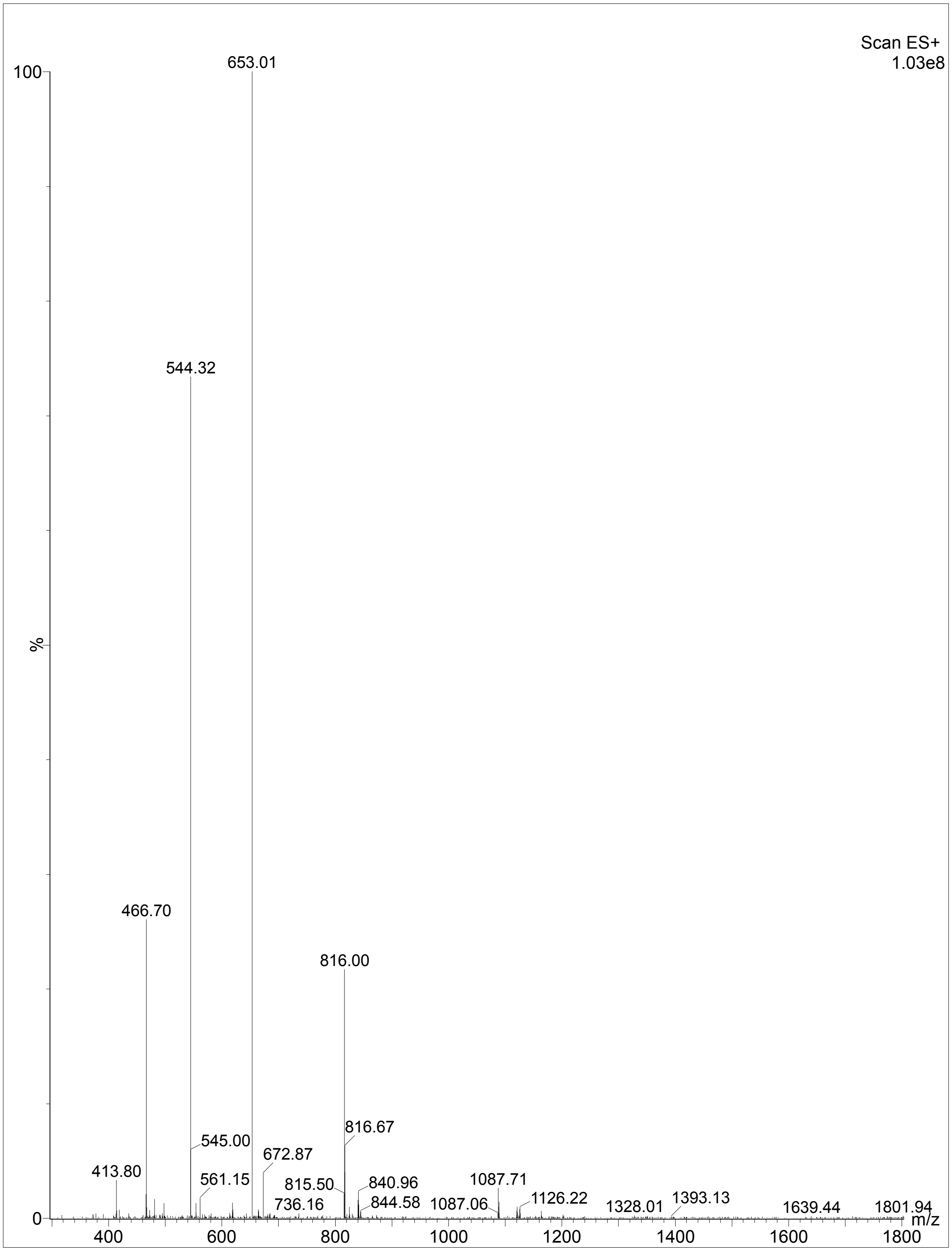


**Figure S32.** Mass spectrometry of P036.

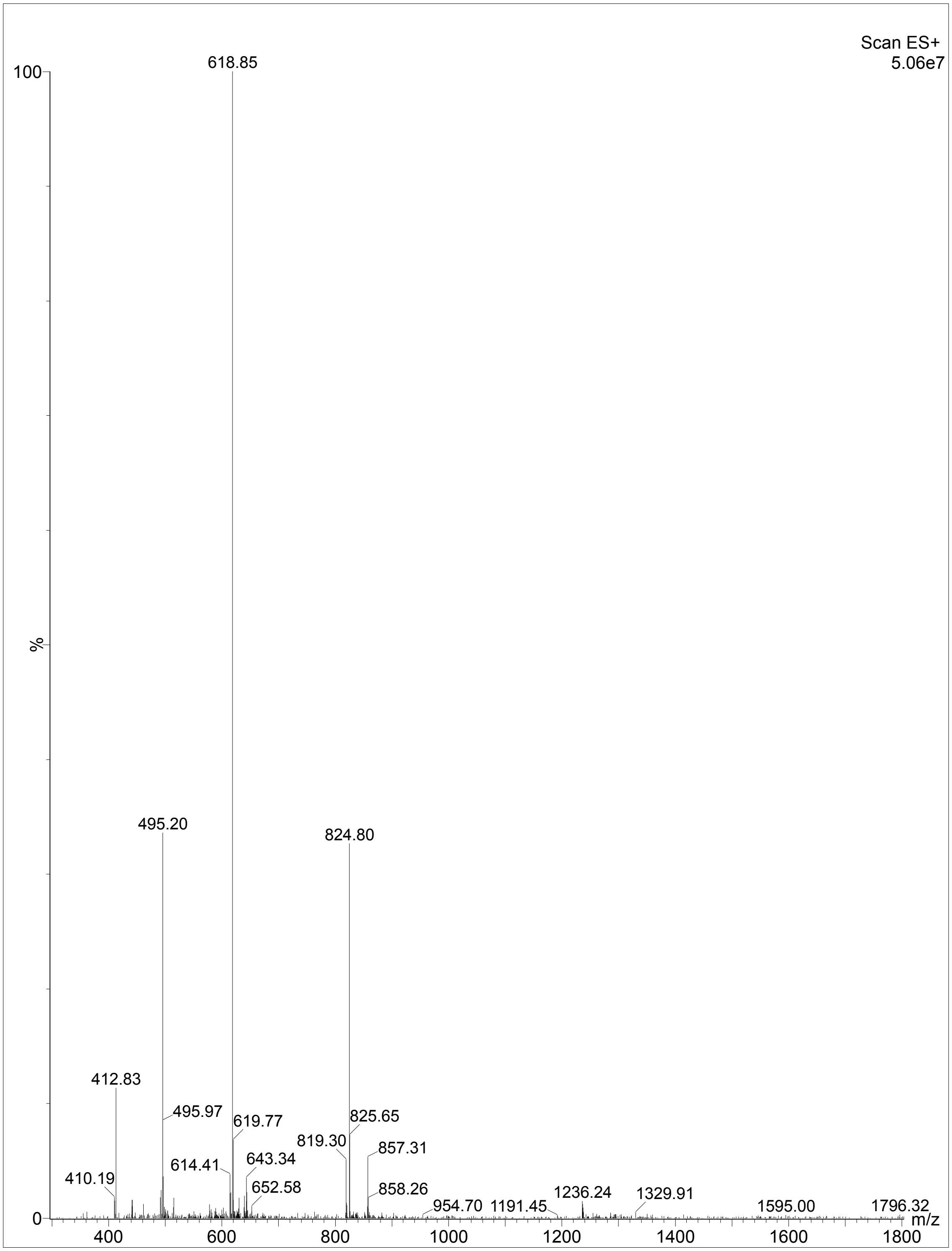


**Figure S33.** Mass spectrometry of P018.

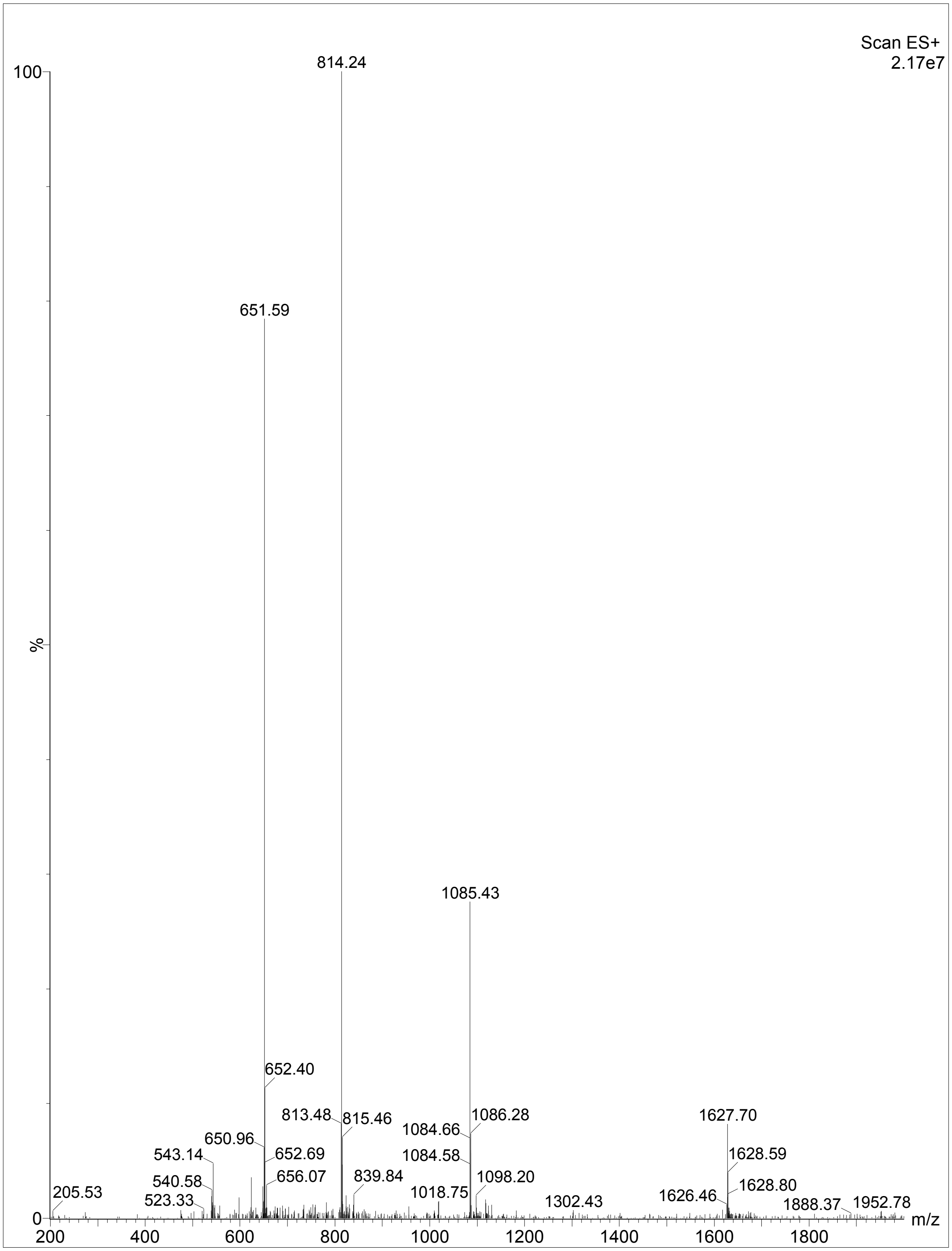


**Figure S34.** Mass spectrometry of P168.

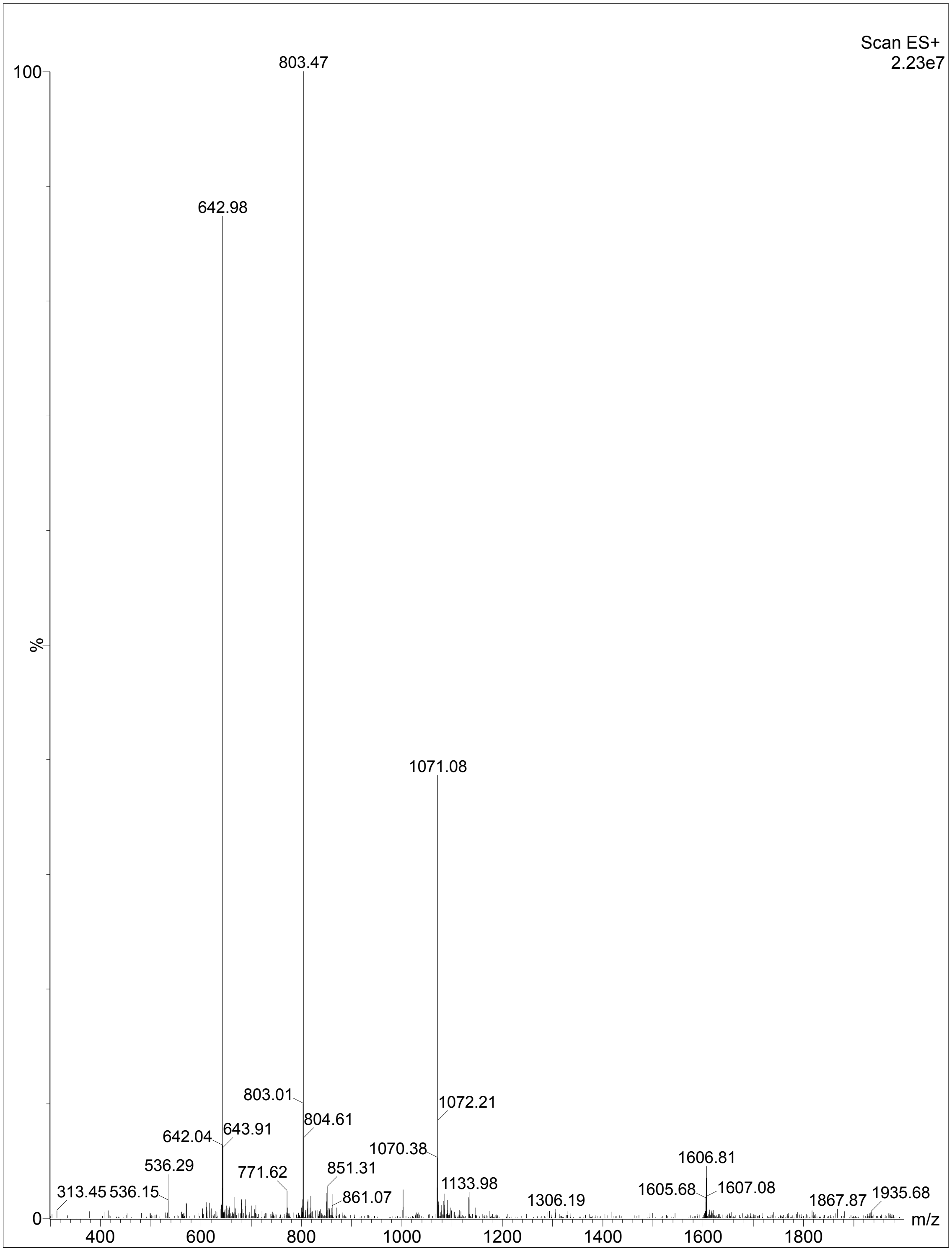


**Figure S35.** Mass spectrometry of P214.

#### Supplementary Tables

**Table S1.** Ablation study results of AMPredictor. Best metrics are marked in bold and second-best values are underlined.

| Fingerprint | Contact Map | ESM | RMSE | MSE | Pearson | CI |
|---|---|---|---|---|---|---|
| √ | × | × | 0.6431 | 0.4126 | 0.4965 | 0.6399 |
| × | √ | × | 0.7273 | 0.5290 | 0.1958 | 0.5406 |
| × | × | √ | 0.5386 | 0.2901 | 0.7032 | 0.6906 |
| √ | √ | × | 0.6543 | 0.4281 | 0.4768 | 0.6540 |
| √ | × | √ | 0.5356 | 0.2869 | 0.6961 | 0.6942 |
| × | √ | √ | 0.5636 | 0.3176 | 0.6494 | 0.7177 |
| √ | √ | √ | **0.5348** | **0.2860** | **0.7072** | **0.7294** |

**Table S2.** Information about five used antiviral classifiers.

| No. | Name | Encoding | Model |
|---|---|---|---|
| 1 | AVPpred | Motif, alignment, AAC, AAindex | SVM |
| 2 | DeepAVP | One-hot | CNN-LSTM |
| 3 | Deep-AVPpred | Pretrained language model | CNN |
| 4 | ENNAVIA | AAC, modlAMP, AAindex | MLP |
| 5 | AI4AVP | PC6 | CNN |

**Table S3.** Sequences and novelty (BLAST E-value) of validated peptides.

| Name | Sequence | Predicted MIC | Length | E-value [a] | E-value [b] |
|---|---|---|---|---|---|
| P089 | GLKALMKILALFMAKKIT | 1.644 | 18 | No hits | 4.1 |
| P386 | PIGLKLRAAIAKGLLHALLKGARAD | 1.857 | 25 | 7.5 | 7 |
| P120 | RWKKILGKAGKLLALRSGKLIL | 1.161 | 22 | 2.6 | 2.4 |
| P135 | KWKQFLKKLAKLIATRIAIIIKRRLK | 1.078 | 26 | 0.85 | 5.3 |
| P247 | KKLGKPLFKGLKKILKLFVKM | 1.450 | 21 | 5.3 | No hits |
| P026 | GIHKKLGPIAKKLLKKIAKL | 3.200 | 20 | 2.7 | No hits |

| | | | | | |
|---|---|---|---|---|---|
| P039 | GWKPILKKAAKIGKRLTGWLFLNSRVN | 2.270 | 27 | 0.29 | 1.2 |
| P070 | LKWWKKFGRKLARKFRM | 2.410 | 17 | 6.6 | No hits |
| P091 | GRMIRKMFKKLWKILRD | 2.480 | 17 | 5.3 | 4.7 |
| P127 | QKPKKDCGGNLLGMIKKFLK | 3.112 | 20 | No hits | 2.7 |
| P122 | IKARLKLKQRIKLIKIGKTFRRRRTDNM | 2.663 | 28 | No hits | 6.9 |
| P352 | RKFKYKFSKLKRVIFMAK | 3.141 | 18 | No hits | 9.9 |
| P105 | FTLKKLLKKGKKTRKLLTPLIKDHSDAM | 1.855 | 28 | No hits | No hits |
| P316 | MFKKILKKATKVIAGLTGHLFWGTRL | 1.317 | 26 | 0.58 | No hits |
| P244 | GTCKGLLKKLLKGMAKFILK | 1.766 | 20 | 6.8 | No hits |
| P302 | GKWQGLLKMIAKILAK | 1.973 | 16 | 0.37 | 0.58 |
| P185 | FRGRIGKILLKMLAQLAK | 2.103 | 18 | 2.3 | 8.3 |
| P019 | PRLAKLLKGGKYILKKITM | 3.482 | 19 | No hits | 0.048 |
| P252 | GMGSFRKLIKKLAKW | 3.491 | 15 | 2.4 | 2.1 |
| P213 | HKCYKWFIKMLRKFQ | 2.982 | 15 | 7.2 | No hits |
| P036 | KMKTFKKKFLKQFAKRLAGRIYPRFK | 2.325 | 26 | No hits | 4.7 |
| P018 | NWKPKIYRTAKKILKIAGKTL | 2.695 | 21 | 2.5 | 2.1 |
| P168 | KKKKFKKFSKGFQKTLMGMLCMSAAKAW | 2.321 | 28 | No hits | 9.1 |
| P214 | KQKIFQKLAARGILLPLAPFIMERIKW | 2.802 | 27 | No hits | No hits |
| P089 | GLKALMKILALFMAKKIT | 1.644 | 18 | No hits | 4.1 |
| P386 | PIGLKLRAAIAKGLLHALLKGARAD | 1.857 | 25 | 7.5 | 7 |
| P120 | RWKKILGKAGKLLALRSGKLIL | 1.161 | 22 | 2.6 | 2.4 |
| P135 | KWKQFLKKLAKLIATRIAIIIKRRLK | 1.078 | 26 | 0.85 | 5.3 |

[a]: E-value was from alignment with the training set.

[b]: E-value was from alignment with DRAMP.

**Table S4.** The $EC_{50}$ (μM) of AMPs inhibiting four enveloped viruses.

| AMP | Generator | CHIKV | HTNV | DENV-2 | HSV-1 |
|---|---|---|---|---|---|
| P001 | v1 | 1.62 | 2.36 | 5.15 | 2.35 |
| P002 | v1 | 0.37 | 2.08 | 0.99 | 0.93 |
| P076 | v1 | 2.03 | 2.26 | 4.48 | 2.73 |
| P135 | v1 | 6.70 | 0.97 | 0.09 | 0.28 |
| P244 | v2 | 42.08 | 2.64 | 1.96 | 2.30 |

**Table S5.** The selectivity index (SI) of three AMPs inhibiting four enveloped viruses.

| AMP | CHIKV | HTNV | DENV-2 | HSV-1 |
|---|---|---|---|---|
| P001 | 37.27 | **50.36** | 11.71 | 25.72 |
| P002 | **149.21** | 31.75 | **51.05** | **59.01** |
| P076 | 22.64 | 15.68 | 14.90 | 16.90 |

**Table S6.** Reported and predicted MICs of some recently mined AMPs.

| No. | Peptide | Sequence | Ref. | Reported MIC | Predicted MIC |
|---|---|---|---|---|---|
| 1 | BigDynorphin | YGGFLRRIRPKLKWDNQKRYGGFLRRQFKVVT | [2] | 14.58 | 7.89 |
| 2 | Apelin-1 | ERPVNLTMRRKLRKHNCLQRRCMPLHSRVPFP | [2] | 41.67 | 13.87 |
| 3 | Apelin-36 | LVQPRGSRNGPGPWQGGRRKFRRQRPRLSHKGPNPF | [2] | 29.17 | 7.75 |
| 4 | vWF-PQR19 | PQRMSRNFVRYVQGLKKKK | [2] | 63.50 | 29.52 |
| 5 | INTb-FTR26 | FTRGKLMSSLHLKRYYGRILHYLKAK | [2] | 25.00 | 6.28 |
| 6 | HSA-GKA38 | GKASSAKQRLKCASLQKFGERAFKAWAVARLSQRFPKA | [2] | 48.92 | 27.66 |
| 7 | CXC19-VKE44 | VKELIKKWEKQVSQKKKQKNGKKHQKKKVLKVRKSQRSRQKKTT | [2] | 76.00 | 0.76 |
| 8 | FIBg-TWK25 | TWKTRWYSMKKTTMKIIPFNRLTIG | [2] | 76.00 | 3.79 |
| 9 | SCUB1-MPF22 | MFPRSFIKLLRSKVSRFLRPYK | [2] | 60.37 | 4.43 |
| 10 | SCUB1-SKE25 | SKEMFPRSFIKLLRSKVSRFLRPYK | [2] | 22.92 | 4.61 |
| 11 | SCUB3-KHK26 | KHKEMLPKSFIKLLRSKVSSFLRPYK | [2] | 20.83 | 5.87 |
| 12 | SCUB3-MLP22 | MLPKSFIKLLRSKVSSFLRPYK | [2] | 18.23 | 3.36 |
| 13 | SFRP1-KKI32 | KKIVPKKKKPLKLGPIKKKDLKKLVLYLKNGA | [2] | 16.67 | 7.41 |
| 14 | NAPP-LIR38 | LIRIPLHRVQPGRRILNLLRGWREPAELPKLGAPSPGD | [2] | 55.17 | 6.14 |
| 15 | NAPP-LIR23 | LIRIPLHRVQPGRRILNLLRGWR | [2] | 87.42 | 20.82 |
| 16 | Bomidin | MGRFKRFRKKFKKLFKKLS | [3] | 4.00 | 5.43 |
| 17 | CBPZ-GSK24 | GSKPWWWSYFTSLSTHRPRWLLKY | [4] | 3.00 | 10.41 |
| 18 | XDH-AVA32 | AVAKLPAQKTEVFRGVLEQLRWFAGKQVKSVA | [4] | 32.00 | 23.51 |
| 19 | c_AMP660 | IFFRRNKKMAVKVAINGFGRIGRLAFRQMF | [5] | 10.00 | 14.56 |
| 20 | c_AMP575 | GRYIAKINPDNKKFKTMPSGKKRKGHKMATHKRKKRLRKNRHKKK | [5] | 2.00 | 1.49 |
| 21 | c_AMP1043 | KQKTLKKVWKLSEKVLIFASAFAKKAGAAEATLVL | [5] | 10.00 | 10.02 |
| 22 | c_AMP67 | AMTLRKRKFAWYVLSSSLKWLIKKAKKIGVQVCGFE | [5] | 20.00 | 4.80 |
| 23 | c_AMP69 | AMTSRKRKFVWYVLSSSLKWLIKKAKKIGVQVCGFE | [5] | 10.00 | 4.66 |
| 24 | c_AMP2041 | SVIWRKLFFIFIKRSGNWIKKVEKRQNLL | [5] | 20.00 | 48.78 |
| 25 | c_AMP250 | DRDRPECSTMVKYEQKLPSLGKYALKRAIKIKFGRK | [5] | 10.00 | 4.40 |
| 26 | c_AMP518 | GINLKRKGNIMKKVKNIFHKIANADPMIWGYVMLSESK | [5] | 25.00 | 6.29 |
| 27 | c_AMP593 | GVPMGSVIKKRRKRMAKKKHRKLLRKTRHQRRNKK | [5] | 25.00 | 2.16 |
| 28 | c_AMP1655 | RGTCYNRVGLIIRNFSKLKGKKV | [5] | 20.00 | 13.73 |

**Table S7.** Minimal inhibitory concentrations (μM) of P076-$NH_2$ and P076.

| AMP | *S. aureus* | | *A. baumannii* | |
|---|---|---|---|---|
| | ATCC 25923 | MRSA | ATCC 17978 | MDRAB |
| P076-$NH_2$ | 53.12 | 106.32 | 53.12 | 53.12 |
| P076 | 6.64 | 13.29 | 0.21 | 0.21 |

**Table S8.** The qRT-PCR primers.

| Virus | Gene | Primers (5'-3') |
|---|---|---|
| HTNV | S | Forward: GAGCCTGGAGACCATCTG<br>Reverse: CGGGACGACAAAGGATGT |
| CHIKV | E | Forward: TCTATAACATGGACTACCCGCCC<br>Reverse: AGCCAGATGGTGCCTGAGAGT |
| HSV-1 | VP16 | Forward: AATGTGGTTTAGCTCCCGCA<br>Reverse: CCAGTTGGCGTGTCTGTTTC |
| DENV-2 | NS5a | Forward: GGTTTTGGGAGCTGGTTGAC<br>Reverse: ACTCTAAGAAGCGTGCTCCA |

**Table S9.** System details of molecular dynamics simulations.

| No. | Peptide | Membrane | Box dimension (nm) | Atoms |
|---|---|---|---|---|
| 1 | P001 | Virus (POPC:POPE:POPS:POPI:DSM :cholesterol = 10:4:1:2:1:2) | 8.66 * 8.66 * 11.60 | 81,660 |
| 2 | P002 | | 8.66 * 8.66 * 11.41 | 80,720 |
| 3 | P076 | | 8.66 * 8.66 * 11.61 | 81,940 |
| 4 | P001 | Inner membranes of G- bacteria (POPE:POPG:TOCL1 = 7:2:1) | 8.13 * 8.13 * 11.60 | 73,644 |
| 5 | P002 | | 8.13 * 8.13 * 11.41 | 72,702 |
| 6 | P076 | | 8.13 * 8.13 * 11.61 | 73,721 |
| 7 | P001 | Outer membranes of G- bacteria (outer: 35 ECLIPA; inner: 75 PPPE, 20 PVPG, 5 PVCL2) | 8.05 * 8.05 * 12.54 | 84,979 |
| 8 | P002 | | 8.15 * 8.15 * 11.88 | 82,483 |
| 9 | P076 | | 8.07 * 8.07 * 12.46 | 84,922 |

**Table S10.** Binding free energy between peptides and the lipid bilayers.

| Peptide | Membrane | $\Delta G_{polar}$ | $\Delta G_{nonpolar}$ | $\Delta G_{bind}$ |
|---|---|---|---|---|
| P001 | Virus | -193.00 ± 16.11 | -44.52 ± 2.99 | -229.65 ± 14.35 |
| P002 | Virus | -136.94 ± 22.66 | -53.60 ± 5.12 | -190.54 ± 18.02 |
| P076 | Virus | -195.71 ± 8.81 | -34.98 ± 8.73 | -230.69 ± 8.31 |
| P001 | G- inner membrane | -408.93 ± 35.39 | -47.97 ± 10.82 | -456.89 ± 29.24 |
| P002 | G- inner membrane | -289.58 ± 49.75 | -70.68 ± 31.58 | -360.26 ± 19.00 |
| P076 | G- inner membrane | -404.77 ± 2.89 | -61.02 ± 10.90 | -465.79 ± 11.39 |
